# Antibody feedback limits specific B-cell entry and alters germinal center repertoires by antibody-antigen complexes

**DOI:** 10.64898/2026.07.29.741558

**Authors:** Keisuke Tonouchi, Joel Finney, Emmanuel James San, Elizabeth Van Itallie, Joshua S. Martin Beem, Dongmei Liao, Xiaoe Liang, Ling Yuan, Scott Szafranski, Masayuki Kuraoka, Kevin R. McCarthy, Kevin Wiehe, Stephen C. Harrison, Garnett Kelsoe

## Abstract

Antibody (Ab) feedback impacts germinal center (GC) responses. Mice given IgG1 or IgG2c recombinant Ab (rAb) specific for the receptor binding site (RBS) of H3 hemagglutinin (HA) and later immunized with the H3 HA trimer exhibited altered GC repertoires. Passive RBS rAb had no effect on the magnitude of ensuing primary GC responses but reduced the numbers of RBS-specific GC B cells. These losses were matched by increases in “unspecific” B cells which did not bind the HA immunogen, with no changes in B cells specific for distal epitopes. These effects were independent of IgG subclass. Higher doses of passive rAb resulted in reduced epitope-specific affinity maturation and clonal proliferation in GCs. Passive rAb generated Ab:HA complexes and favored recruitment of rare HA-specific GC B cells with enhanced avidity for the rAb:HA immune complex (IC). We show that some no- and low-affinity GC B cells represent responses to local ICs.

## Introduction

Antibody (Ab) is the principal effector arm of humoral immunity acting systemically to protect against pathogens^1^. Ab-secreting plasmacytes are often products of germinal center (GC) responses in which somatic B-cell evolution generates cells expressing higher-affinity B-cell antigen receptors (BCR). These selected cells, on their differentiation into plasmacytes, are the sources of avid and protective serum Ab^2^.

Among the factors affecting the quality and magnitude of humoral responses are the Ab products of the response itself^3–5^. Currently, it is thought that a major effect of circulating Ab in secondary humoral responses is the masking of epitopes that dominated earlier humoral responses. This Ab masking would then interfere with *de novo* recruitment of B cells specific for those same epitopes or with the persistence of these cells in secondary GCs^6–10^. A significant longer-term effect of this epitope-masking is thought to be the broadening of the humoral response to otherwise sub-dominant epitopes^8,10^.

In addition to epitope-masking, a significant literature suggests a role for the isotype of circulating Ab in the modulation of secondary humoral responses^5,11–13^. B cells express the inhibitory FcγR IIb, whose signals raise the threshold of BCR activation^14,15^. Reciprocally, the CD21/CD35 complement receptors lower that activation threshold in cooperation with stimulatory coreceptors on B cells^16–19^. Therefore, mouse IgG1 Ab, which cannot fix complement and binds most avidly to FcγR IIb (CD32), is thought to inhibit secondary responses when complexed with antigen, whereas IgG2a/c Ab may enhance them, by complement fixation to bound antigen^3^. In addition to direct effects on B cells, Ab/antigen immune complexes (ICs) promote antigen retention by C3-decoration of ICs on follicular dendritic cells (FDC) through CD21/CD35, binding IC to FDC by CD32, or by interacting with FcγRs on myeloid antigen-presenting cells^13,20,21^.

The “end-product inhibition” of adaptive humoral responses has been long recognized as a critical factor in the quality of Ab-mediated immunity elicited by infection or vaccination^4^. Recent studies and reviews have highlighted the correlation between pre-existing Ab and post-boost B-cell and Ab repertoires elicited in humans by repeated administration of SARS-CoV-2 or influenza vaccines^22–28^. These studies could not, however, discriminate the effects of circulating Ab from other immune factors (*e.g*., elicited Tfh repertoires^29,30^). Because that these potentially confounding actors are absent in primary humoral responses, several recent studies have also examined the effects of passive transfer of serum or recombinant Ab(s) (rAb) and/or the use of BCR knock-in mouse models^7,8,10^. Although studies with BCR knock-in animals show reduced recruitment of specific BCR knock-in cells to GCs, the artificially restricted clonality poorly represents physiological BCR heterogeneity and selection. Moreover, no prior study has investigated the effects of Ab feedback on the “unspecific” GC B cell populations which have no readily detectable specificity for native immunogen but constitute ≥40% of primary GC responses^31^.

Here, we measured the effects of Ab feedback on physiological primary GC responses elicited by HA immunization. The magnitude of elicited GC responses was the same whether mice received passive rAb specific for the HA receptor binding site (RBS), rAb to an irrelevant antigen, or in untreated (No Ab) controls. In contrast, frequencies of HA-specific GC B cells were about 35% lower in mice given RBS Ab than in the No Ab or irrelevant Ab control groups; this loss was the result of significant reductions in the GC B cells specific for the RBS and adjacent (RBS/interface) epitopic sites. Responses to other epitopic sites were unaffected both in rank order of immunodominance and in proportion to total GC B-cell responses.

Losses of RBS and RBS/interface GC B cells were not compensated by clonal increases/expansions of B cells targeting alternative epitopic sites but rather by increased numbers of “unspecific” cells that did not bind to the HA immunogen. In all three experimental groups, BCRs mapped to specific epitopic sites showed substantial genetic homogeneity, except for the BCRs specific for the RBS and RBS/Interface in mice given passive RBS rAb. In that group, the reduced populations of GC B cells specific for RBS and RBS/Interface had a strong bias (44% of mice) for a stereotyped IGHV14-4/IGKV4-59 receptor, which were present but rare in control animals (<1%). High-resolution structural analyses of the corresponding IGHV14-4/IGKV4-59 Fabs bound with a complex of HA and the Fab of the passive RBS Ab revealed that their epitopes included a contribution of the transferred RBS Ab, as well as from the HA surface: the presence of passive RBS Ab presumably advantages this particular response. This large data set clarifies the effects and mechanisms of Ab feedback and provides a novel explanation for the recruitment of “unspecific” B cells into persistent GC responses.

## Results

### Experimental outline

To determine the independent effects of circulating Ab on the B-cell repertoires of primary GC responses, we transferred low (20μg) or high (200μg) doses of the musinized human K03.12 Ab^32^ (HA RBS specific) or mouse BMPC23 rAb^33^ (HSV gB specific) by *i.v.* injection, followed 48 hours later by immunization with H3 A/Kansas/14/2017 X-327 HA trimer (H3/KS) in alum (Figure 1A). Prior work has shown that influenza infection or HA vaccination elicit dominant Ab responses to the HA RBS and nearby epitopic sites^34–36^; the epitope defined by the K03.12 Ab falls within the H3/KS RBS (K_D_ = 72nM) (Figure S1A). The BMPC23 Ab is specific for the gB glycoprotein of HSV and has no detectable reactivity for H3/KS even at 200µg/ml (Figure S1B). To control for any unspecific effects of the passive IgG1 or IgG2c rAbs, a second age-matched control group (No Ab) was immunized without rAb pretreatment.

**Figure 1.**
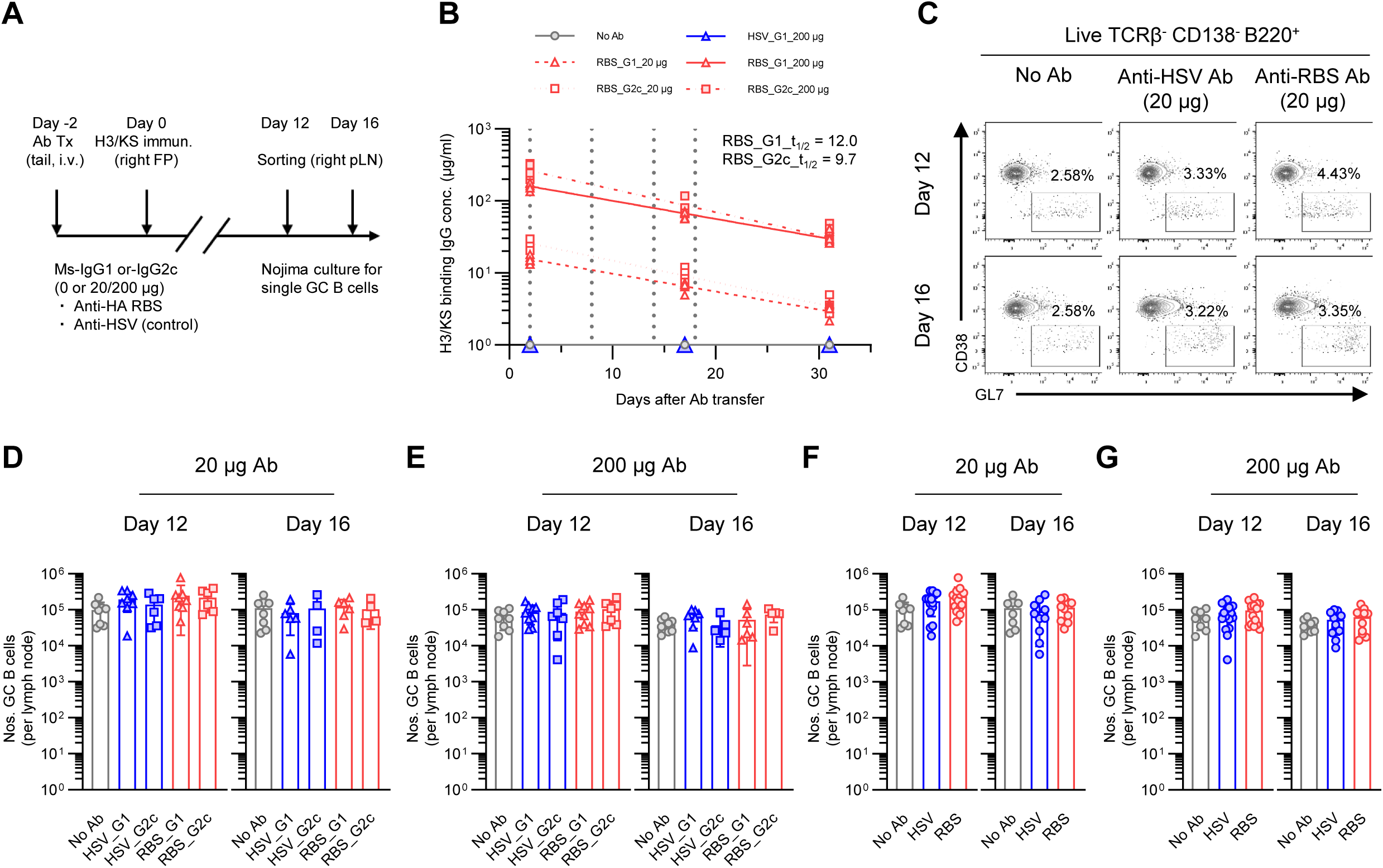
No effect of passive Ab on the magnitude of GC responses. (A) A scheme of the experiment. (B) Kinetics of passive Abs in mice after the transfer. Serum IgG concentration specific to H3/KS was measured in wild-type C57BL/6 given 20µg of musinized K03.12 IgG1 (RBS_G1_20µg, red clear triangle with dashed line) or -IgG2c (RBS_G2c_20µg, red clear square with dotted line), or 200µg of same K03.12 IgG1 (RBS_G1_200µg, red filled triangle with solid line) or -IgG2c (RBS_G2c_200µg, red filled square with dashed/dotted line). Mice infused with 200µg of BMPC23 IgG1 (HSV_G1_200µg, blue filled triangle with solid line) or without any Ab (No Ab, gray filled circle with solid line) as controls. Each plot represents individual mice (n=5 biological replicates from single experiment). Values were collectively fitted by nonlinear curves using GraphPad Prism. The days corresponding to Ab transfer, HA immunization, d12- and d16 sampling in later experiments are highlighted with dotted lines. Half-lives of passive RBS IgG1 and IgG2c mAbs are indicated on right top. (C) Representative flow plots for GC B cell gating (CD38^dull^ GL7^+^) in mice transferred with No Ab (left), and 20µg of HSV Ab (middle) or RBS Ab (right) on days 12 (upper) and 16 (lower). Live TCRβ^-^ CD138^-^ B220^+^ population were gated. The frequencies of GC B cells among the total B cells are indicated above each gate. (D-G) Calculated numbers of GC B cells per collected pLN are shown by (D and E) separating groups transferred with IgG1 (triangle) and IgG2c (square) or (F and G) pooling those IgG subclasses (circle). Each plot represents individual animals from No Ab (gray), HSV Ab (blue), or RBS Ab (red) cohorts with (D and F) 20µg or (E and G) 200µg of passive transfer. The data are pooled from 4-8 experiments (n=4-8 biological replicates without combining IgG subclasses). Bars indicate mean±SD. *P*-values were calculated by Kruskal-Wallis test with Dunn’s multiple comparisons: no statistical significance was seen. See also Figure S1.

The decay rates for passive musinized K03.12 were determined in unimmunized mice to have similar half-lives (t_½_) for IgG1 and IgG2c at t_½_=12d and 9.7d, respectively (Figure 1B). We estimated the concentrations of passive rAb in blood and lymph to have been 88% of the initial transfer amount on the day of immunization (day 0), 60% at the onset (day 6), 41% at peak (day 12), and 32% at subsiding stages (day 16) of GC responses to HA/alum^31^ (Figure 1B).

GC B-cell responses were sampled on days 12 and 16 in single-cell Nojima cultures^31,37^. Flow cytometric enumeration and isolation of individual GC B cells was based on their CD38^dull^ GL7^+^ surface phenotype^31,37^. At harvest, individual Nojima cultures were initially screened for IgG production and then antigen- and epitope-specificities and avidities were determined in validated Luminex assays^31,32,37,38^. BCR V(D)J rearrangements of HA-reactive GC B clones were subsequently recovered and analyzed^37^.

### No effect of passive Ab on the magnitude of GC responses

Flow cytometric enumeration of GC B cells in mice receiving the passive RBS Ab, HSV Ab and No Ab groups showed comparable and robust GC responses at days 12 and -16 (Figure 1C). Passive Ab had no significant effect on the numbers of GC B cell elicited or on the frequencies of GC phenotype B cells in the draining lymph node (LN) (Figures 1D-1G, S1C-S1F). We could detect no significant differences in the effects of passive IgG1 or IgG2c Abs at either tested dose (Figures 1D, 1E, S1C and S1D). Thus, the capacity of passive IgG2c RBS Ab to fix complement and interact with activating FcγR had no measurable effect on the magnitude or kinetics of primary GC responses compared to passive IgG1. Consequently, for the rest of our study we combined the results for both isotypes when comparing the effects of passive RBS and HSV Abs (Figures 1F, 1G, S1E and S1F). Disaggregated data for the two isotypes are provided in relevant supplemental figures. We conclude that supraphysiologic levels of circulating Ab with good affinity for a major epitope present on the immunogen had no effect on the magnitude or kinetics primary GC responses.

### Passive RBS Ab skews GC recruitment of antigen-specific and -unspecific B cells

To determine whether passive RBS Ab altered the repertoire of responding GC B-cell populations, from 129 mice we established 184,412 single-cell Nojima cultures^31^ for GC B cells and recovered 33,536 (18% cloning efficiency) IgG^+^ clonal cultures for analysis of the specificity and genetics of GC B cells in each experimental cohort (range, 1,895 - 4,845 cells) (Table. S1).

Culture supernatants were screened for IgG, κ-, and λ-L chain concentrations and binding to H3/KS as described^31^. A total of 10,771 IgG^+^ clonal cultures reacted with H3/KS (Table. S1). In both control groups, HA-binding cultures accounted for 33-39% and 31-54% of IgG^+^ cultures on days 12 and -16, respectively, regardless of passive Ab dose (Figure 2). In mice receiving 200 µg of passive RBS Ab, frequencies of HA-binding cultures were significantly lower on both day 12 and -16 and those given 20 µg RBS Ab showed significantly reduced numbers of HA-specific clones on day 12 but not day 16 (Figures 2A and 2C). Thus, whereas the HA^+^ GC compartment was reduced by passive RBS Ab, the magnitude of GC responses was not (Figures 1F and 1G). Recruitment of “unspecific” or “dark antigen”^31^ GC B cells was increased by the presence of passive RBS Ab (Figures 2B and 2D). The effects of passive IgG1 and IgG2c RBS Ab were indistinguishable (Figure S2).

**Figure 2.**
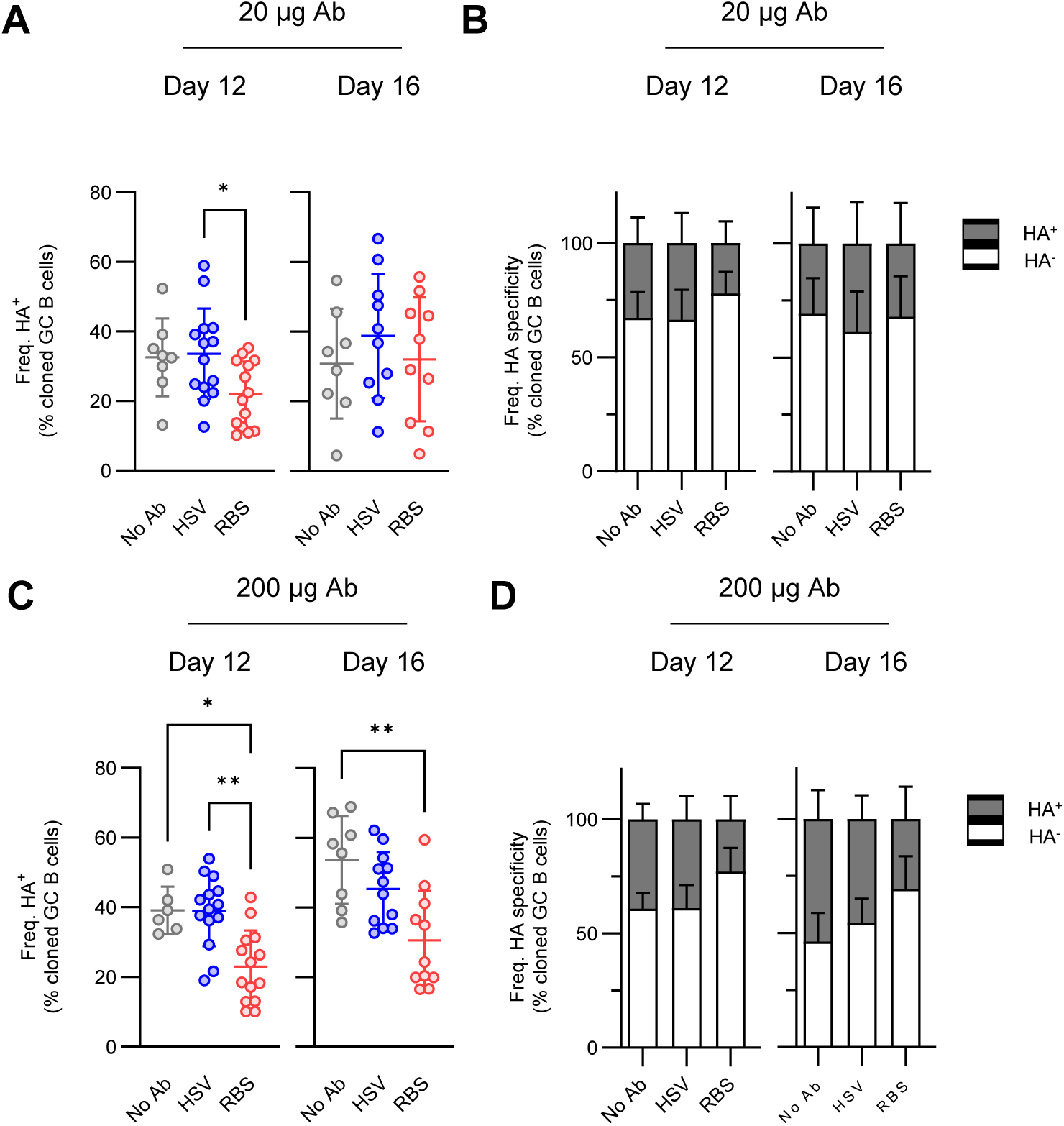
Passive RBS Ab skews GC recruitment of antigen-specific and -unspecific B cells. (A and C) The frequency of H3/KS reactive GC B cell cultures among total IgG secreting clonal cultures. Each plot represents individual animals from No Ab (gray), HSV Ab (blue), or RBS Ab (red) cohorts with (A) 20µg or (C) 200µg of passive transfer. Bars indicate mean±SD. *P*-values were calculated by Kruskal-Wallis test with Dunn’s multiple comparisons: \**p* < 0.05, \*\**p* < 0.01. (B and D) The ratio of H3KS specific (gray) and -unspecific (white) GC B cell cultures among total IgG secreting clonal cultures. Bars indicate mean+SD of HA^+^ or HA^-^ population in each group. (A-D) The data are pooled from 4-8 experiments (n=6-14 biological replicates with combining IgG subclasses). See also Figure S2 and Table S1.

### Repertoire selection for epitopic sites other than RBS and RBS-interface is unchanged by the presence of passive RBS Ab

To determine whether passive RBS Ab reduces the frequency of H3/KS-binding GC B cells in an epitope-specific fashion, we mapped the epitopes of H3/KS-binders using a panel of mutant H3/KS proteins^39,40^ that identify binders to six canonical epitopic sites^32,38,41–47^ (Figures 3A and S3A). These mutations reduce (> 80%) binding of well-characterized rAbs known to be specific for each epitopic site, *e.g*., RBS^32,41,45^, head interface (Interface)^48^, vestigial esterase domain (Esterase)^46^, *etc*., without impacting the binding of standard Abs directed to other epitopes (Figure 3B).

**Figure 3.**
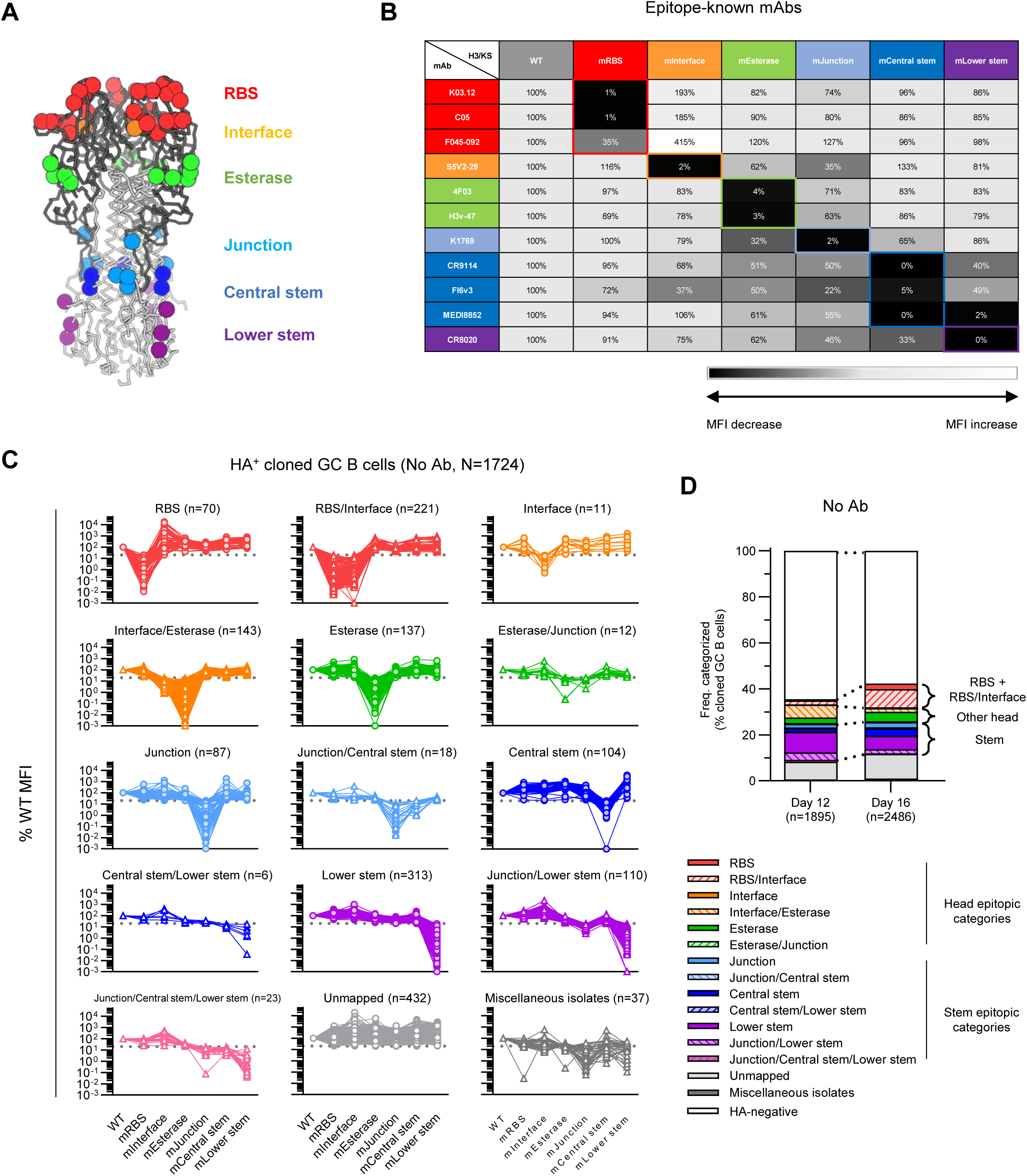
Epitope mapping identifies the diversity and the shift of clonal selection in GC induced by complex antigen. (A) Structural diagram of H3 HA. The HA structure of A/Aichi/02/1968, X-31 (H3N2) (PDB: 2VIU)^87^ is utilized. Locations of mutations introduced in each H3/KS mutants are mapped by spheres, which are colored by epitopes corresponding to six canonical sites (RBS, red; Interface, orange; Esterase, green; Junction, light blue; Central stem, dark blue; Lower stem, purple). HA head domain is shown in dark gray, and stem domain is shown in light gray. Side view of the molecule is indicated. (B) Validation of the H3/KS mutant panel using epitope-known mAbs. The percentages of MFI values relative to the WT H3/KS construct are shown. Each row represents the values in individual mAbs against each column of H3/KS constructs. The values are color-coded with indicating stronger reduction of binding in darker colors (0% in black). The colors of mAbs and H3/KS construct names are matched by their epitope specificity and the corresponding mutations described in (A). The specific reductions based on the known-epitope specificity are highlighted in colored rectangles. (C) Evaluation of the H3/KS mutant panel using HA^+^ GC B cell cultures. The percentages of MFI values relative to the WT H3/KS construct are shown. Each plot represents individual cultures, and values from the same sample are connected with a line. HA specific cultures of No Ab group obtained in Figure 2 (N=1,724, pooling all the day 12 and day 16 samples) were divided into 15 epitopic categories, and each category of GC B cell cultures is shown separately with the number of corresponding cultures on the top of individual figures. Specific reduction was set by 80% reduction and the threshold line (20% of WT MFI) is indicated by dotted lines. The cultures which were mapped by two or more mutants but not assigned by any of 13 mapped categories were pooled and defined as miscellaneous isolates (dark gray). (D) Frequency distribution of 15 epitopic categories among total IgG^+^ GC B cell cultures. Data from the samples in (C) are shown along with the proportion of HA^-^ cultures (white) with dividing day of the sampling. The numbers of cultures are indicated at the bottom of each column. The frequencies were calculated by individual animals (n=14-16 biological replicates) and subdivided bars represent the mean of them. Grouping of some epitopic categories (RBS+RBS/Interface, other head, and stem) are indicated with connecting day 12 and day 16 fractions by dotted lines. See also Figure S3 and Table S2.

In this way, we characterized epitope specificity of clonal IgGs from individual Nojima cultures, considering any mutant HA that reduced IgG binding by >80% (relative to the wildtype H3/KS) to indicate epitopic specificity (Figures 3C, S3B and S3C). Some clonal IgGs had reduced binding to adjacent mutated sites, *i.e*., these IgGs mapped to epitopes spanning two or three mutated sites. Along with the six canonical epitopic categories, we defined 7 additional ones; RBS/Interface, Interface/Esterase, Esterase/Junction, Junction/Central stem, Central stem/Lower stem, Junction/Lower stem, and Junction/Central stem/Lower stem. Clonal cultures specific for canonical (48%, 5,120/10,771) and intermediate (26%, 2,831/10,771) epitopic categories comprised 74% of all HA^+^ GC B cell cultures (Table S2). About 25% of clonal cultures (2,707/10,771) showed no loss of binding to any H3/KS mutant (Figures 3C, S3B and S3C, and Table S2), *i.e*., they bound known epitopes not identified by the mutant panel (*e.g*., post-fusion epitopes)^39,49,50^ or at novel epitopic sites. Clonal IgGs recognizing these unmapped epitopes were present in all mice studied (129/129), indicating that these unmapped HA epitopes are robustly immunogenic in mice. Finally, rare clonal IgGs (1%) had reduced binding to ≥2 mutants not included in the seven intermediate categories defined above. These rare clone types were observed only occasionally (range, <1% - 9% of tested mice) in all cohorts. We pooled these infrequent IgGs and categorized them as miscellaneous isolates (Figures 3C, S3B and S3C, and Table S2).

Absent passive Ab (No Ab), individual HA^+^ Nojima cultures mapped specifically to one of the 15 epitopic categories with clear hierarchy of immunodominance (Figure 3D). On day 12 after immunization, RBS-mapped cultures (RBS+RBS/Interface) accounted for 2±2% of all clonal IgG cultures, while other head categories constituted 8±4% and pooled stem categories made up 16±6%. By day 16, the frequency of RBS+RBS/Interface clones increased to 10±9%, with other head and stem epitopes at 6±5% and 14±7%, respectively. Unmapped HA^+^ clones also increased over this interval, from 8±6% on day 12 to 11±6% on day 16. This method of mapping single GC B cells to individual epitopic sites permits detailed characterization of GC repertoire diversity and dynamics during primary responses to a complex protein antigen^31^.

All experimental cohorts showed significant variability among the individual age-matched female mice within single experiments (Figure S4A-S4D). This variability was unrelated to HA or other reagent lots. We suggest this variability represents an intrinsic property of B6 mice responding to a complex T-dependent protein Ag.

In both the No Ab and HSV Ab groups, distributions of epitope specificity were similar on both days 12 and -16 regardless of passive Ab dose (Figures 4A and 4B). In the RBS Ab group at day 12, both low and high Ab doses resulted in modestly lower frequencies for RBS+RBS/Interface cultures (0.5±0.9% and 0.6±1.2%, respectively) than for controls (≈2%) (Figures 4C and 4D), but by day 16 the effect was stronger, 0.7±0.5% and 0.3±0.4% (Figures 4C and 4D) vs 4-16% in controls. Results were equivalent for IgG1 and IgG2c RBS rAb groups (Figure S4E and S4F).

**Figure 4.**
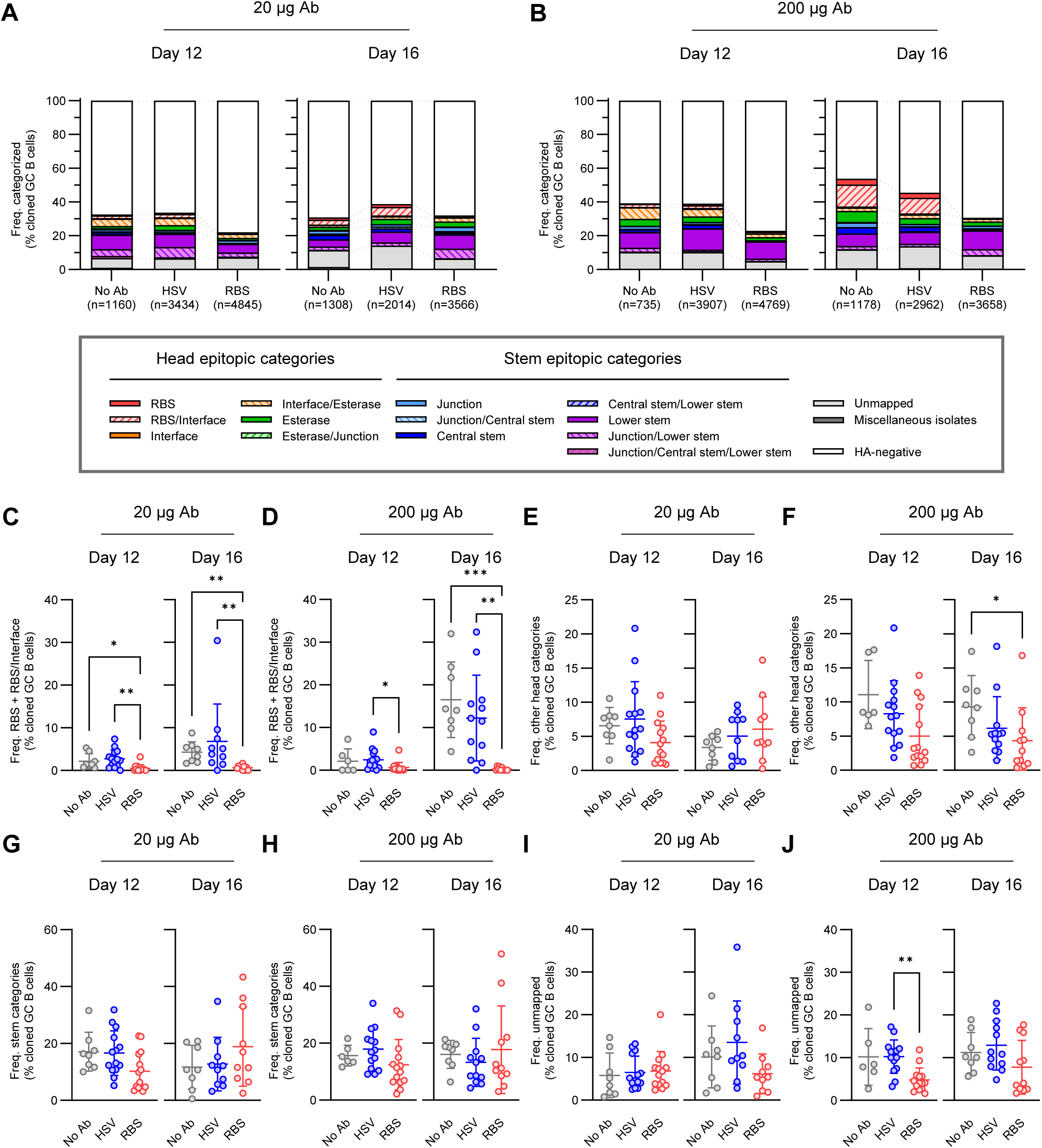
Repertoire selection for epitopic sites other than RBS and RBS-interface is unchanged by the presence of passive RBS Ab. (A and B) Frequency distribution of 15 epitopic categories among total IgG^+^ GC B cell cultures. HA specific clonal cultures obtained in Figure 2 were examined for their epitope specificities. The numbers of IgG^+^ cultures, including HA^-^ cultures, are indicated at the bottom of each column. No Ab, HSV Ab, or RBS Ab group with (A) 20µg or (B) 200µg of passive transfer are shown. The frequencies were calculated by individual animals (n=6-14 biological replicates with combining IgG subclasses) and subdivided bars represent the mean of them. Grouping of some epitopic categories (RBS+RBS/Interface, other head, and stem) are indicated with connecting each group of fractions by dotted lines. (C-J) Frequency of GC B cell cultures mapped as (C and D) RBS and RBS/Interface, (E and F) other head, or (G and H) stem, or defined as (I and J) unmapped among total cloned population. Each plot represents individual animals from No Ab (gray), HSV Ab (blue), or RBS Ab (red) cohorts with (C, E, G and I) 20µg or (D, F, H and J) 200µg of passive transfer (n=6-14 biological replicates with combining IgG subclasses). Bars indicate mean±SD. *P*-values were calculated by Kruskal-Wallis test with Dunn’s multiple comparisons: \**p* < 0.05, \*\**p* < 0.01, \*\*\**p* < 0.001. See also Figure S4 and Table S2.

While passive RBS Ab reduced the frequency of RBS+RBS/Interface clones (Figures 4C and 4D), GC selection for other HA epitopic sites was not enhanced (Figures 4E-4J). This observation was true for both day 12 and -16. The stronger effects on RBS-binders at day 16 had no discernable effect on the immunodominance rank order of other epitopic sites. Passive RBS Ab feedback did not shift responses to distal antigenic determinants.

### Suppression of affinity maturation to the RBS epitopic site by passive RBS Ab

To determine if Ab feedback affects affinity maturation, we determined the avidity index (AvIn)^31^ relative to K03.12^32^ (K_D_ = 72nM) (Figure S1) for each HA-reactive culture. We necessarily excluded from analysis the few H3/KS-binding cultures (<7%) with binding MFI values too low for comparison to K03.12. We determined AvIn values for 10,040 GC B-cell clonal cultures (Figures 5A, 5B, S5A and S5B). Consistent with earlier reports, AvIn values were broadly distributed (≈10^-5^ – ≈10^0^) in all cohorts both on day 12 and day 16, consistent with permissive selection in GC B cell populations^31,38,51^. At both low and high doses, neither passive HSV Ab nor RBS Ab had any general effect on affinity maturation in primary GCs (Figures 5A and 5B). The absence of any general effect on affinity maturation was true regardless of the IgG subtype of passive Ab (Figures S5A and S5B). In all control and experimental groups, we observed little change in median AvIn values between days 12 and 16 (Figures 5A and 5B), suggesting that any substantial affinity maturation had taken place before the peak GC response at day 12^31^.

**Figure 5.**
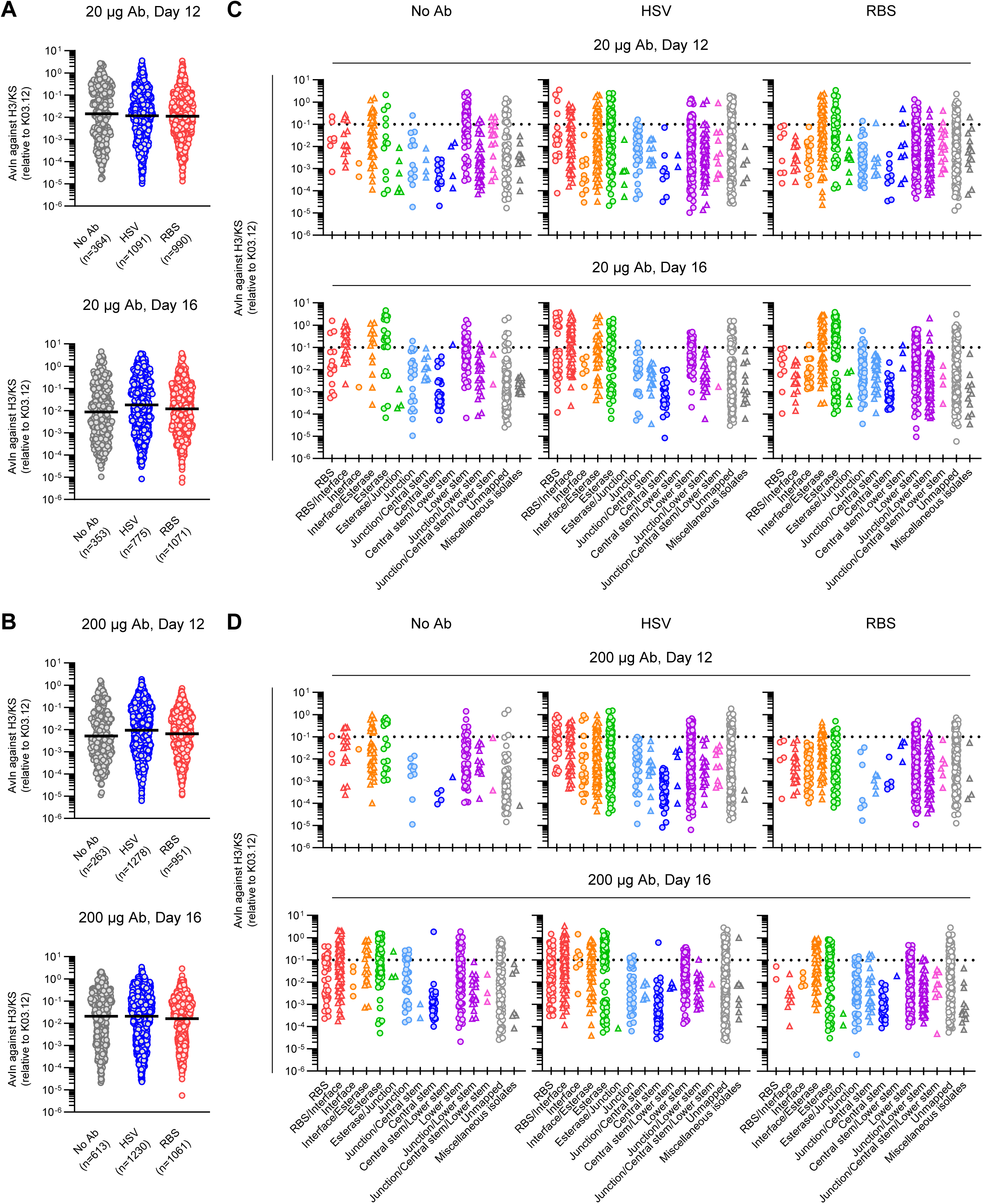
Suppression of affinity maturation to the RBS epitopic site by passive RBS Ab. (A and B) Distribution of avidities in H3/KS specific GC B cell cultures. HA specific clonal cultures obtained in Figure 2 were analyzed. The numbers of samples which were able to calculate AvIn are indicated at the bottom of each column. Each plot represents individual GC B cells from No Ab (gray), HSV Ab (blue), or RBS Ab (red) cohorts with (A) 20µg or (B) 200µg of passive transfer. Bars represent median. (C and D) AvIn distribution obtained in (A) and (B) were broken by epitope specificity. Data under the condition of (C) 20µg or (D) 200µg of passive transfer are shown. Cutoff lines for high avidity GC B cells (AvIn > 0.1) are indicated by dotted lines. See also Figure S5.

While passive RBS Ab had little or no impact on affinity maturation of the majority of HA specific GC B cells, it affected the abundance and affinity maturation of GC B cells mapped to RBS and RBS/interface epitopic sites (Figures 5C and 5D). We observed significantly fewer RBS and RBS/Interface in the passive RBS Ab group than in controls (except for the anomalously low day 12 population of RBS+RBS-interface Abs in the No Ab group), with the loss of high-avidity compartments (AvIn values ≥0.1) (Figures 5C, 5D, S5C and S5D). Passive RBS Ab had no advantage on the clonal abundance and AvIn distributions of GC B cells targeting non-RBS epitopic sites (Figures S5E-S5J).

### Reduced clonal expansion and cell proliferation at supraphysiological doses of passive RBS Ab

We expected epitopic site-specific competition by the high-affinity K03.12 Ab to favor recruitment of higher-avidity RBS-binding B cells to GCs. Instead, higher-avidity cells were specifically censored, especially on day 16 (Figures S5C and S5D). To investigate whether the residual, less-avid clones (n=88) were subject to the same GC selection as others, we sequenced the V(D)J gene rearrangements present in all H3/KS^+^ GC B cell cultures (n=10,771) and recovered 6,919 complete H- and L-chain V(D)J sequences. This allowed us to characterize mutation frequencies and clonal sizes of GC B cells in each experimental cohort (451-806 cells of sequences/cohort) (Table. S3).

Consistent with the absence of a general effect on affinity maturation (Figure 5), H3/KS-reactive GC B cells from all cohorts accumulated comparable frequencies of nucleotide mutations in V_H_ genes (Figures 6A and 6B). Accrual of additional mutations between days 12 and 16 did not correspond to an increase in median avidity for H3/KS (Figures 5A and 5B), implying that GC B cell proliferation during that interval supported the incorporations of net-neutral mutations with respect to BCR avidity for H3/KS.

**Figure 6.**
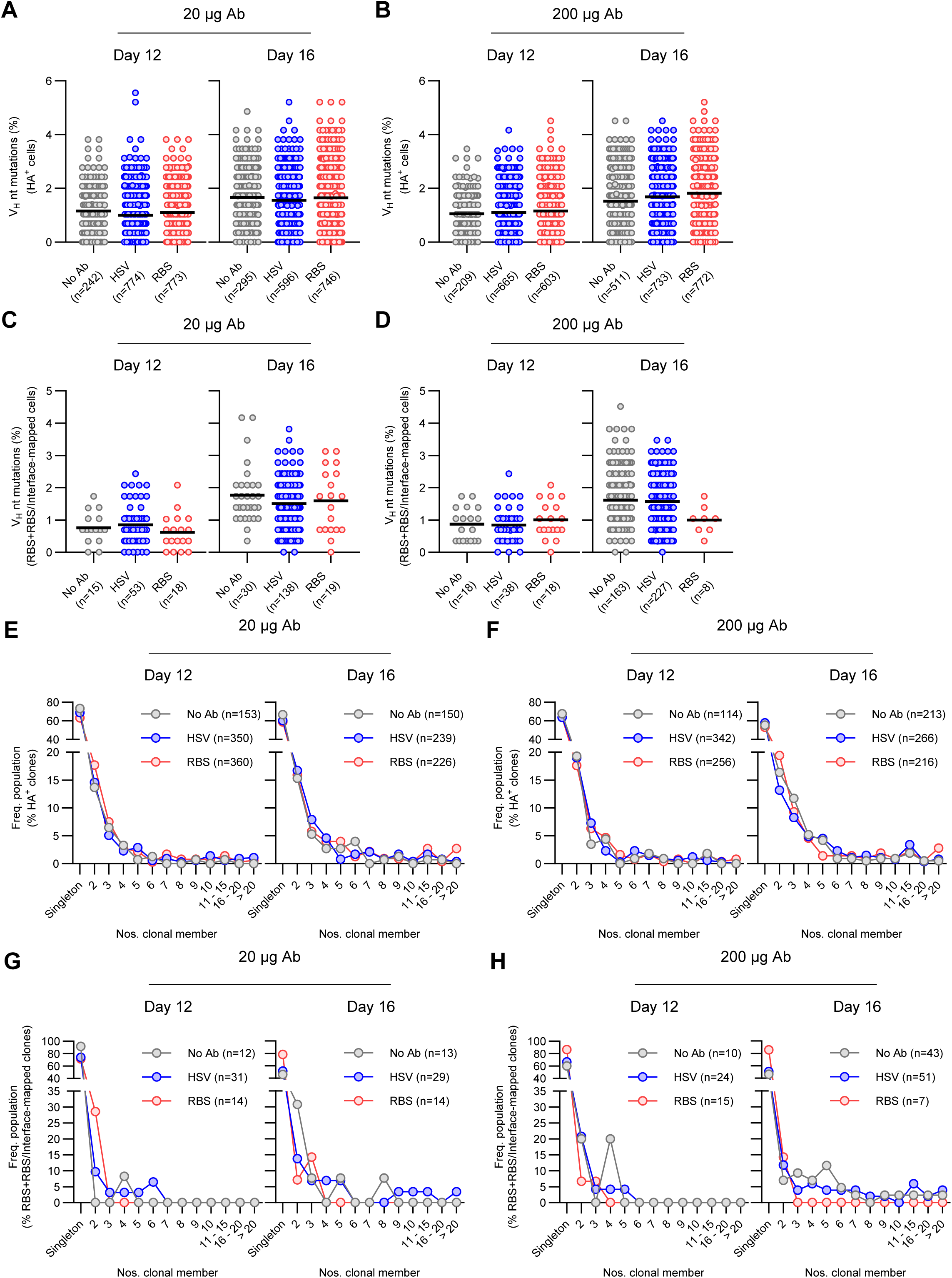
Reduced clonal expansion and cell proliferation at supraphysiological doses of passive RBS Ab. (A and B) Distribution of V_H_ mutation frequency in H3/KS reactive GC B cell cultures. HA specific clonal cultures obtained in Figure 2 were sequenced. The numbers of samples which were successfully recovered with complete heavy plus light chain pairs are indicated at the bottom of each column. Each plot represents individual sequenced GC B cells from No Ab (gray), HSV Ab (blue), or RBS Ab (red) cohorts with (A) 20µg or (B) 200µg of passive transfer. Bars represent mean. (C and D) Among the sequences obtained in (A) and (B), distributions of V_H_ mutation frequency in GC B cell cultures mapped as RBS and RBS/Interface are shown. The numbers of corresponding cells are indicated at the bottom of each column. Each plot represents individual sequenced GC B cells from No Ab (gray), HSV Ab (blue), or RBS Ab (red) cohorts with (C) 20µg or (D) 200µg of passive transfer. Bars represent mean. (E and F) Frequency distribution of clone size in H3/KS reactive GC B cell cultures. Sequences obtained in (A) and (B) were analyzed for their clonal lineage, and the numbers of the clonal members in individual unique clones were enumerated. The numbers of the unique clones identified are indicated at the right top of each figure, and the frequencies of each clone size were calculated by that number. Histograms for No Ab (gray), HSV Ab (blue), or RBS Ab (red) cohorts with (E) 20µg or (F) 200µg of passive transfer are shown. (G and H) Among the unique clones obtained in (E) and (F), frequency distribution of clone size in GC B cell clones mapped as RBS and RBS/Interface are shown. The numbers of corresponding clones are indicated at the right top of each figure, and the frequencies of each clone size were calculated by that number. Histograms for No Ab (gray), HSV Ab (blue), or RBS Ab (red) cohorts with (G) 20µg or (H) 200µg of passive transfer are shown. See also Figure S6.

Numbers of V_H_ mutations in GC B cells specific for the RBS+RBS/Interface from mice in the low-dose RBS Ab cohort did not differ from GC B cells from control animals (Figure 6C): mean(±SD) frequencies of V_H_ mutations were comparable to those in total HA^+^ cells. RBS+RBS/Interface GC B cells from mice in the high-dose RBS Ab cohort, however, had typical frequencies (1±1%) of V_H_ mutations on day 12 but, unlike controls, had accrued no more V_H_ mutations by day 16 (Figure 6D). We could not determine the statistical significance of this difference due to the low numbers of GC B cells recovered from mice in the RBS Ab groups. The IgG subclass of passive Ab made no difference in the accumulation of mutations (Figures S6A-S6D).

Lower mutation frequencies in RBS+RBS/Interface GC B cells from RBS Ab-treated mice than from controls might result from restricted proliferation in the absence of available epitopes to drive BCR activation. If so, clonal expansion would be restricted only in GC B-cell clones competing with passive Ab. To test this hypothesis, we compared clone sizes in the passive RBS and control groups. Cloanalyst^52^ identified 2,885 unique clones among the sequenced H3/KS reactive GC B cells (Table. S3). In both low and high dose passive transfers, the distribution of the clone size was identical among No Ab, HSV Ab and RBS Ab cohorts at both days 12 and 16 (Figures 6E and 6F). On day 12, >87% of the clones in each group had no more than 1-3 members; large clones with ≥10 members accounted for <5% of all HA^+^ cultures. By day 16, the frequency of large clones increased, but small clone sizes of 1-3 members remained dominant in all experimental groups (>79% of total). Thus, passive RBS Ab did not generally affect clonal size. We found no evidence for clonal bursts in GC B cells elicited by H3/KS, consistent with a recent report^53^.

In clones mapped as RBS+RBS/Interface, clone sizes of 1-3 members predominated in all treatment groups (63-100% of total) (Figures 6G and 6H). The No Ab and HSV Ab controls did, however, have substantial numbers of GC B clones with ≥4 clonal members on day 12 (8-20%) and day 16 (15-37%), while no such clones were recovered from mice given either dose of RBS Ab. Moreover, the frequency of singletons was higher in mice receiving 200μg of RBS Ab than in those given 20μg of RBS Ab, both on day 12 (87% vs 71%) and day 16 (86% vs 79%) (Figures 6G and 6H). Even at the lower dose tested, passive RBS Ab limited clonal expansion of GC B cells specific for the RBS and RBS/Interface epitopic sites. The higher dose of passive RBS Ab had a stronger effect; it presumably suppressed cell proliferation, leading to the reduced accumulation of mutations we observed.

### No effect of passive RBS Ab on the BCR genetics of cells mapped to non-RBS+RBS/Interface epitopes

Even though passive K03.12 significantly inhibited the selection of RBS-specific GC B, selection for other HA reactive cells was unchanged. To examine whether there was an effect on the clonal composition of these populations specific for other, distal epitopic regions, we compared the use of V_H_ and V_L_ gene segments in clonal cultures not mapped to RBS and RBS/Interface epitopic sites (other head, stem, unmapped) and compared the HSV Ab and No Ab controls to the RBS Ab group.

A substantial majority of all mouse V_H_ gene segments (84 of 110)^54^ were recovered from control and experimental groups on day 12 and -16 (Figure S7). Despite this genetic diversity, each epitopic category showed genetic preferences. In all three experimental groups, IGHV1-53 and IGHV9-3 were common in clones mapped as other head; IGHV1-50 and IGHV1-69, in clones mapped as stem; and IGHV1-66 and IGHV1-81 in unmapped clones. These preferences did not change between days 12 and -16, with the exception of higher representation for IGHV3-6 in stem-targeted clones on day 16. We suggest that inter-clonal selection of GC B-cell repertoires was largely complete by day 12 postimmunization. These preferences for V_H_ gene segments were unchanged by passive RBS Ab (Figure S7).

We conclude that the Ab feedback effect against GC selection was rigorously specific to epitopic site and the reduction of the competed B cells did not significantly alter the selection of other epitope-specific BCR repertoires.

### Improved recruitment of rare B cells in the presence of passive RBS Ab by cooperative enhancement of affinity for HA

The presence of low-affinity GC B cells specific for the RBS+RBS/Interface epitopic sites in the presence of high-affinity passive RBS Ab is not consistent with a process of direct competition at the binding site. Instead, it might result from infrequent, chance selection or by recruitment of B cells specific for a non-RBS determinant disrupted in the RBS and/or Interface HA mutants. To determine the similarity of RBS+RBS/Interface GC B cells from RBS Ab mice to those from No Ab and HSV Ab controls, we compared V_H_ gene use in each group. In both control groups, BCRs of GC B clones mapped as RBS+RBS/Interface were strongly enriched for the IGHV1-47 gene segment (average 45% and 58% on days 12 and 16, respectively) (Figures 7A and 7B). In contrast, IGHV1-47 usage was low (7%) in mice receiving the low dose of RBS Ab and was absent in animals receiving the high dose (Figures 7A and 7B). Instead, mice given passive RBS Ab were significantly enriched for the IGHV14-4 segment (21% - 86%) in clones mapped to the RBS+RBS/Interface sites (Figures 7A and 7B).

**Figure 7.**
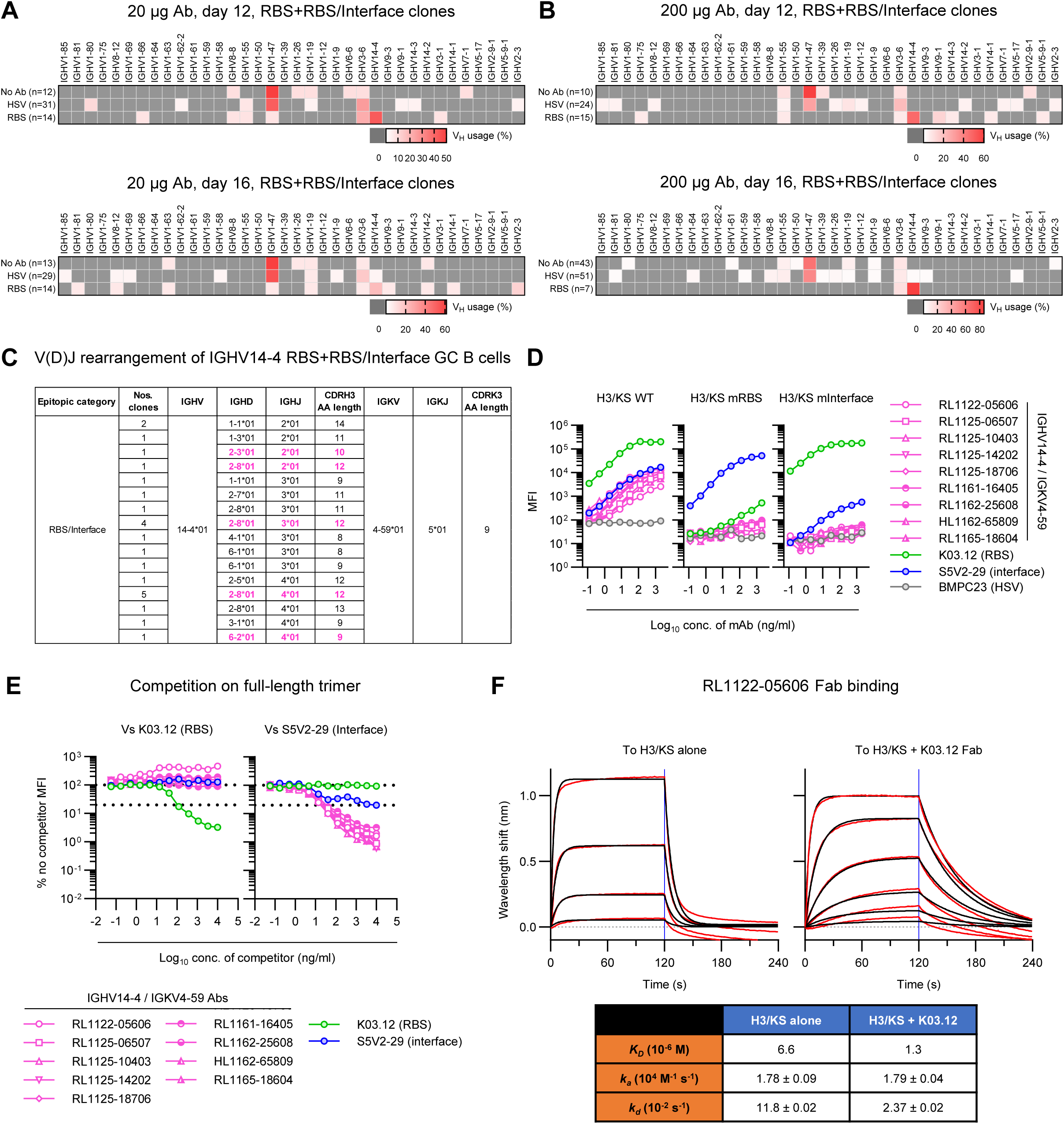
Improved recruitment of rare B cells in the presence of passive RBS Ab by cooperative enhancement of affinity for HA. (A and B) V_H_ gene usage of cultured GC B cells mapped as RBS and RBS/Interface. Heatmaps for No Ab, HSV Ab, or RBS Ab cohorts with (A) 20µg or (B) 200µg of passive transfer are shown. The numbers of unique clones mapped as RBS+RBS/Interface are indicated next to the names of groups, and the frequencies of each V_H_ gene usage were calculated by that number. Values are color-coded with higher frequency in thicker red (0% in dark gray). All the V_H_ genes identified in RBS+RBS/Interface mapped GC B cells throughout the experiments are shown in the order encoded in the gene locus (left, distal to constant region; right, proximal to constant region). (C) V(D)J usage of IGHV14-4 GC B cells identified within RBS+RBS/Interface clones. The table shows the epitope specificity, number(s) of unique clone(s), V(D)J usage and CDR3 length in amino acid of heavy and light chains. D_H_/J_H_ genes and HCDR3 length of clones which were utilized by creating mAbs in later figures are highlighted in pink color. (D) Binding curve of IGHV14-4/IGKV4-59 mAbs against WT-, RBS mutant-, and Interface mutant H3/KS constructs. Each mAb including controls (K03.12, RBS mAb; S5V2-29, Interface mAb; BMPC23, HSV mAb as negative control) was serially diluted from 2µg/ml and assessed in an Luminex assay. MFI values at each dilution are plotted. (E) Competition assays of IGHV14-4/IGKV4-59 mAbs against RBS or Interface mAb. Binding of IGHV14-4/IGKV4-59 mAbs were assessed on H3/KS in the presence of K03.12 (RBS) and S5V2-29 (Interface). BMPC23 (HSV) was utilized as a negative control competitor and considered as no competitor condition. Competitor mAbs were serially diluted from 10µg/ml and applied ahead of adding sample mAbs. Percentage of MFI values relative to no competitor at each dilution are plotted. The dotted lines indicate 100% (upper) and 20% (lower) of no competitor MFI. (F) Affinity measurement of K03.12 against H3/KS alone (left) and H3/KS+K03.12 (right). Kinetics of binding intensity was measured using BLI. Measured and fitted curves are shown in black and red, respectively. Blue line segregates the association (left side) and dissociation (right side) curves. Calculated K_D_ value and on/off-rates (k_a_ and k_d_) are shown below the graph.. (D and E) Data are the representative of at least three repeat experiments. See also Figure S7.

IGHV14-4 BCRs were not restricted to the RBS Ab groups: three (0.11%; 3/2,768) clonal GC B cultures recovered from two HSV Ab control mice on day 16 used the IGHV14-4 gene segment to encode BCRs specific for RBS+RBS/Interface epitopic sites (Figures 7A and 7B). These rare examples indicate that passive K03.12 Ab promoted GC entry and persistence of RBS+RBS/Interface reactive B cells expressing IGHV14-4 BCRs, but that such cells could enter GC in its absence.

All IGHV14-4 RBS+RBS/Interface BCRs, in both control and RBS Ab groups, contained a single, stereotyped κ L-chain (IGKV4-59/IGKJ5) and a common KCDR3 sequence: QQWSSNPLT or QQWSSSPLT (Figure 7C). To characterize the epitopic site(s) recognized by these IGHV14-4/IGKV4-59 RBS/Interface BCRs, we generated nine IgG1 rAbs randomly selected from the 32 stereotyped BCRs identified. Consistent with the original epitopic mapping of using culture supernatant IgGs, all nine rAbs bound WT H3/KS but neither RBS-nor Interface-mutant HAs (Figure 7D).

We conducted competition assays for the nine stereotyped RBS/Interface rAbs against the RBS mAb (K03.12) and an Interface rAb, S5V2-29^38^. Homologous inhibition by K03.12 and S5V2-29 produced concentration-dependent inhibition of H3/KS binding (97% and 80%, respectively), but no cross-competition (Figure 7E). S5V2-29 potently inhibited binding of all nine IGHV14-4/IGKV4-59 rAbs to H3/KS (>97% inhibition) but K03.12 inhibited none of them. Binding of one IGHV14-4/IGKV4-59 rAb, RL1122-05606, was enhanced >4-fold in the presence of K03.12 (Figure 7E). Biolayer interferometry (BLI) measurements confirmed that in the absence of K03.12, RL1122-05606 Fab had modest affinity for H3/KS (K_D_ = 6.6µM) (Figure 7F) but when the immobilized antigen was a complex of HA plus K03.12 Fab, the affinity of RL1122-05606 was 5-fold higher (K_D_ = 1.3µM). Reduction in the dissociation rate (from 11.8±0.02 x 10^-2^ s^-1^ to 2.37±0.02 x 10^-2^ s^-1^) accounted for the affinity increase (Figure 7F). Thus, binding of K03.12 to HA directly enhanced the affinity of RL1122-05606 for the HA immunogen.

### IGHV14-4/IGKV4-59 RBS/Interface mAbs bind a complex epitope comprising HA plus K03.12

To identify the epitope of the IGHV14-4/IGKV4-59 rAb, RL1122-05606, from RBS Ab treated mice, we used single-particle cryogenic electron microscopy (cryo-EM) to determine its structure bound to a H3/KS:K03.12 IC. For comparison, we also determined the structure of the IGHV14-4/IGKV4-59 rAb, HL1162-65809, isolated from a control HSV Ab mouse, bound to the H3/KS:K03.12 IC. The preparations contained the H3/KS head, K03.12 Fab, and either RL1122-05606 Fab or HL1162-65809 Fab (Figures S8 and S9). We also included S8V1-157 Fab, which binds a head-stem interface epitope^50^, to aid particle-picking and alignment. The 3-D reconstructions of these complexes had resolutions of 2.9 Å (RL1122-05606) (Figure S8) and 3.2 Å (HL1162-65809) (Figure S9), respectively.

Atomic models fit to the cryo-EM maps showed that both RL1122-05606 and HL1162-65809 bind similar, complex epitopes containing residues from both HA and the K03.12 HC variable domain (Figures 8A, S8F and S9F). The HA portion of the epitopes includes parts of the 220- and 130-loops around the rim of the RBS, plus some of the head interface (the surface on the side of the head where two HA protomers would contact each other) (Figures 8B, S10A and S10B). The epitope:paratope interfaces of RL1122-05606 and HL1162-65809 are generally conserved, but the HCDR3 of HL1162-65809 makes more contacts with HA than does the HCDR3 of RL1122-05606 (Figures S10A and S10B). Those additional contacts probably explain why HL1162-65809 binds more tightly to HA alone (K*_D_* ≍ 1μM) than does RL1122-05606 (K*_D_* ≍ 7μM) (Figures 7F and S10C). RL1122-05606 and HL1162-65809, along with seven other IGHV14-4/IGKV4-59 mAbs from mice given RBS Ab, exhibited no measurable affinity for K03.12 alone (Figure S10D), showing that their main contacts are with HA and that binding to K03.12 is supplemental.

**Figure 8.**
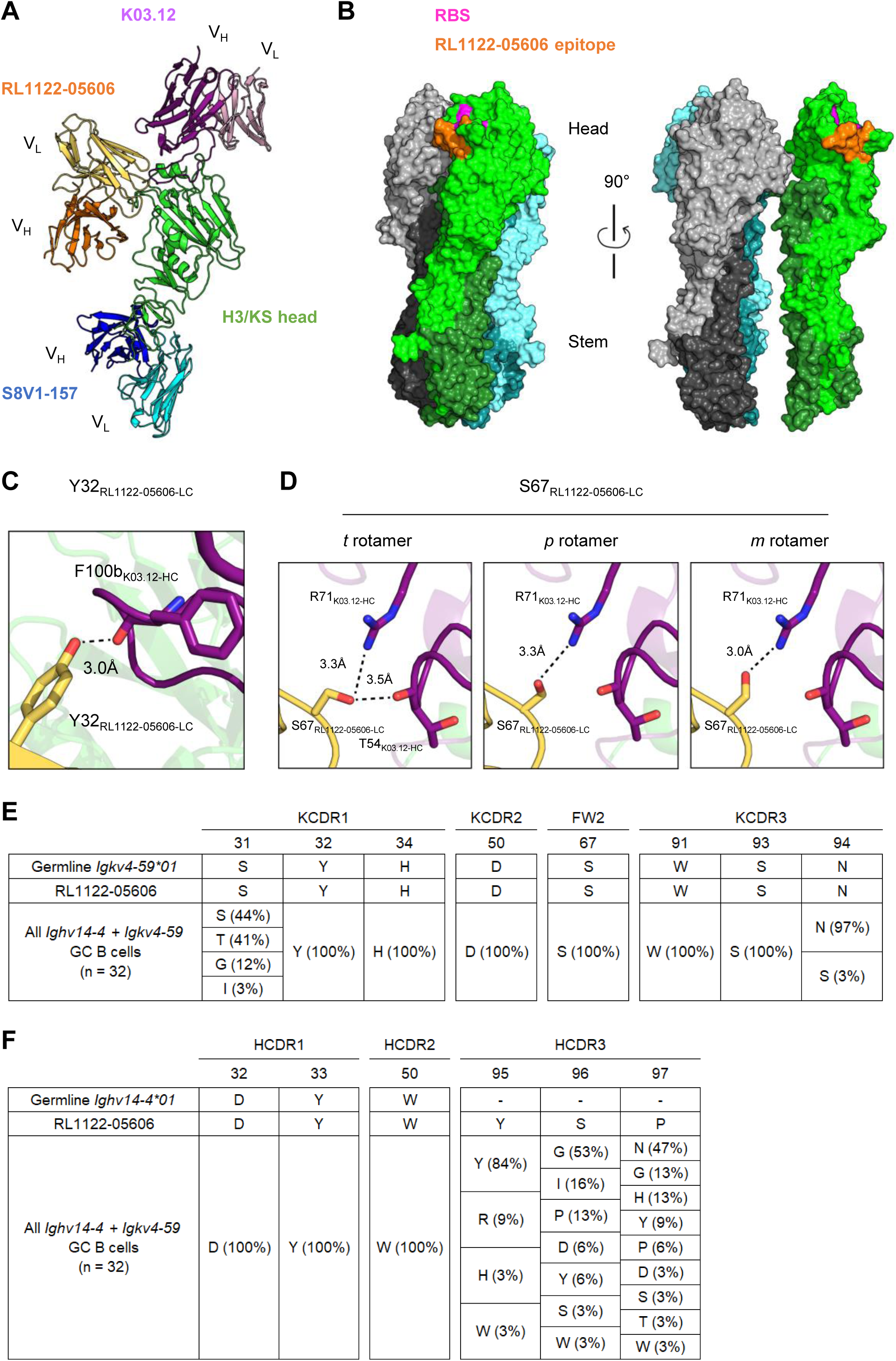
IGHV14-4/IGKV4-59 RBS/Interface mAbs bind a complex epitope comprising HA plus K03.12. (A) A cartoon rendering of an atomic model of the complex of RL1122-05606 Fab (orange), K03.12 Fab (purple), S8V1-157 Fab (blue), and H3/KS head (green), as determined by cryo-EM. (B) Surface rendering of the atomic model of homotrimeric HA ectodomain from influenza A/Aichi/02/1968, X-31 (H3N2) (PDB: 2VIU)^87^. Each HA protomer is colored distinctly, and the HA2 subunits are shaded darker than the HA1 subunits. In the left panel, the green protomer is oriented identically to the HA head shown in (A). The pink and orange regions respectively denote the RBS and RL1122-05606 footprint. (C and D) Close-up views of the hydrogen-bonds (dashed black lines) between the RL1122-05606 LC (yellow) and the K03.12 HC (purple). The three panels in (D) show that the hydroxyl group of S67RL1122-LC binds the K03.12 HC regardless of the sidechain rotamer. (E and F) Alignments of the paratope residues of the RL1122-05606 LC (E) or HC (F) with their inferred germline sequences and with the peptide sequences from all RBS/interface-binding GC B cells that use IGHV14-4/IGKV4-59-encoded BCRs. See also Figures S8-S10.

The K03.12 portion of the epitope includes a hydrogen bond from Y32_LC_ to the backbone carbonyl of F100b_K03.12-HC_ (Figure 8C), and a hydrogen bond between the side-chain of S67_LC_ and N53_K03.12-HC_ and/or R71_K03.12-HC_ (Figure 8D). Y32_LC_ and S67_LC_ are conserved in all IGHV14-4/IGKV4-59 RBS/Interface Abs (Figure 8E); thus, it is likely that all such Abs bind the K03.12 HC, enabling at least one of these Abs bind more tightly to HA:K03.12 complex than to HA alone (Figures 7E and 7F). We note that the hydrogen bonds between HL1162-65809 and K03.12 do not impart higher avidity of HL1162-65809 for HA:K03.12 than for HA alone (Figure 7E).

Restricted BCR diversity is often associated with antigen/epitope specificity in humans, with well-known examples being the influenza HA stem^55,56^ and -head interface^57,58^, SARS-CoV-2 receptor binding domain (RBD)^59^, and HIV-1 envelope protein (Env)^60,61^. In mice, Abs targeting certain occluded HA epitopes also exhibit restricted Ab/BCR diversity^62,63^. The IGHV14-4/IGKV4-59 RBS/Interface Abs bind the composite HA+K03.12 epitope through germline encoded residues that contact K03.12, Y32_LC_ and S67_LC_, which are present in >60 IGKV gene segments deposited in IMGT database. Restriction to IGKV4-59 may be favored in that it uniquely codes for residues D50 and N94, which together accept three hydrogen-bonds from R222_HA_ and R224_HA_ (Figures 8E, S10A and S10B); no other IGKV gene segment can plausibly recapitulate these interactions. Every IGHV14-4/IGKV4-59 Ab/BCR that we examined retains both D50_LC_ and N94_LC_ (Figure 8E), as expected for critical paratope residues. Genetic restriction to IGHV14-4 may derive largely from the side chains of paratope residues 32 and 33. Y33_HC_ makes several nonpolar contacts with HA, and its hydroxyl group also accepts a hydrogen bond that no other side chain would be long enough to reach (Figures 8F, S10A and S10B). Y33 is present in ≍25% of germline IGHV gene segments. D32_HC_ contacts HA with its Cα and/or Cβ atoms; interactions that could be reconstituted by many other side chains (Figures 8F, S10A and S10B). The structures of RL1122-05606 and HL1162-65809 suggest that bulky side chains probably are disfavored at residue 32; five IGHV genes, including IGHV14-4, encode both Y33 and a non-bulky sidechain at residue 32. All IGHV14-4/IGKV4-59 RBS/interface Abs preserve the D32_HC_ and Y33_HC_ residues (Fig 8F).

### IGHV14-4/IGKV4-59 GC B cells that do not bind HA

That the epitope recognized by IGHV14-4/IGKV4-59 mAbs was a composite of HA and RBS Ab immediately suggested that similar GC B cells might be among those initially identified as unspecific (HA^-^). Screening HA^-^ Nojima cultures for paired IGHV4-14 and IGK4-59/JK5 rearrangements showed an additional 102 GC B cells expressing this stereotyped BCR (Table 1). The great majority of these stereotyped GC cells were recovered from mice given passive RBS Ab but 3 examples were recovered from No Ab and HSV Ab mice (Table 1). This single stereotyped BCR represented almost 1.5% (99/6,918) of all HA^-^ GC B-cell cultures from RBS Ab mice.

**Table 1.** Unspecific (HA^-^) GC B cell cultures expressing a IGHV14-4 and IGKV4-59/JK5 BCR.

| Experimental Group |  |  | Nos. mice sequenced | Nos. H+L sequences | GC B cells<br>VH14-4 +<br>Vk4-59/Jk5 | Nos. mice<br>stereotyped<br>BCR |
| --- | --- | --- | --- | --- | --- | --- |
| Day 12 | 20 µg | No Ab | 8 | 453 | 0 | 0 |
|  |  | HSV | 14 | 1519 | 0 | 0 |
|  |  | RBS | 14 | 2620 | 4 | 2 |
|  | 200 µg | No Ab | 4 | 260 | 0 | 0 |
|  |  | HSV | 8 | 1064 | 2 | 2 |
|  |  | RBS | 8 | 1778 | 87 | 5 |
| Day 16 | 20 µg | No Ab | 8 | 593 | 1 | 1 |
|  |  | HSV | 10 | 722 | 0 | 0 |
|  |  | RBS | 10 | 1444 | 1 | 1 |
|  | 200 µg | No Ab | 4 | 223 | 0 | 0 |
|  |  | HSV | 4 | 640 | 0 | 0 |
|  |  | RBS | 6 | 1076 | 7 | 1 |
| Total |  |  | 98 | 12392 | 102 | 12 |

## Discussion

Experiments describing the effects of passive “Ab feedback” were first reported by Uhr and colleagues in the 1960s^64^. Resurgent, contemporary interest in the serological, cellular and molecular analyses of secondary responses to SARS-CoV-2 or influenza vaccines in humans^22–25,27,28^ and mice^65,66^ has refocused attention on the feedback phenomenon. In BCR knock-in mice, passive transfer of primary serum Ig or specific recombinant Ab constrains the repertoire of naïve knock-in B cells recruited by boost immunizations^7,10^. These studies, however, do not exclude a potential role for other immune factors, *e.g*., selective T_REG_ activity, in the regulation of post-boost Ab repertoires. For this reason, we have investigated in C57BL/6 mice the effect of a single, passive Ab (RBS Ab, K03.12) in regulating the selection and maturation of B cells in primary GCs elicited by a complex protein antigen (H3/KS).

We found that supraphysiological amounts of passive RBS Ab had no effect on the magnitude or the kinetics of subsequent primary GC responses elicited by HA immunization, despite significant reductions in the frequencies of GC B cells specific for the adjacent RBS and RBS/Interface epitopic sites. These site-specific reductions had no systematic effect on selection for distal epitope regions but instead were compensated by increases in “unspecific” GC B cells lacking detectable binding for the H3/KS HA immunogen. These observations indicate that passive RBS Ab did not reduce GC responses by antigen sequestration but rather impaired the activation of B cells specific for the RBS and RBS/Interface regions to prevent or limit entry into the GC reaction. We also suggest that the magnitude of GC responses may be regulated by factors other than the numbers or density of B-cell epitopes on antigen.

Passive RBS Ab also suppressed affinity maturation in primary GC B cells recognizing RBS or RBS/Interface epitopic regions but had no effect on those specific for other epitopic sites. Passive RBS Ab did not alter the BCR genetics of GC cells mapped to non-RBS+RBS/Interface epitopes, which remained similar in all experimental cohorts and comprised a broad range of heavy- and light-chain variable gene segments. An IGHV14-4/IGKV4-59 BCR rarely recovered from control mice was more frequent in mice given passive RBS Ab. Representative IGHV14-4/IGKV4-59 rAbs from both control and RBS Ab groups were generated and most bound the IC of H3/KS HA and K03.12 RBS Ab, with higher affinity than for HA alone. High-resolution structures showed that the binding of IGHV14-4/IGKV4-59 rAbs was determined by contributions from both HA and RBS Ab. Passive RBS Ab promoted recruitment of B cells specific for the IC into GCs, not recruitment of B cells recognizing a novel or normally subdominant HA epitope. Our experiments are consistent with the effects of Ab feedback being determined by the specificity and V-regions of passive RBS Ab. The effects of passive IgG1 or IgG2c RBS Ab were indistinguishable whereas complementation of IGHV14-4/IGKV4-59 rAbs depended on interaction with K03.12 RBA Ab V-domains.

Various molecular mechanisms have been suggested to explain the controlling effects of circulating Ab on B-cell responses^3^. Whereas several receptors expressed on B cells have demonstrated inhibitory or enhancing effects on B cell activation^14–19^, whether circulating Ab interacts with these to control GC responses has been controversial. Ab isotype has been suggested to play a role in modulating antigen presentation by dendritic cells, FDCs or subcapsular sinus macrophages either directly via FcγR engagement or indirectly through CD21/35 via complement fixation^13,20,21^. Recent studies have suggested that IgG isotype differences have no or little role in Ab feedback effects^7,67^. Our results show identical effects of passive RBS IgG1 or IgG2c Ab and substantiate these conclusions.

Prior studies have reported inhibited recruitment of antigen-specific naïve or memory B cells to GCs in the presence antigen-specific Ab^3–6,66,68^; recent animal studies have highlighted that this epitope-specific suppression^7,10^ shifts the B cell/serum Ab repertoires to other epitopic sites present in the boost immunogen^8,65,69,70^. Our data confirm epitope-specific inhibition by passive Ab, but we did not observe changes in the rank order or genetic preferences of other GC B-cell specificities. Given that (*i*) we studied Ab inhibition of a single epitope (RBS) and (*ii*) RBS+ RBS/Interface-mapped GC B cells were not initially immunodominant (5%-9% of HA^+^ GC B cultures in controls), we cannot exclude the possibility that suppression of the RBS+RBS/Interface response was insufficient to alter the later immunodominance of other HA determinants.

Passive K03.12 Ab markedly increased the presence of GC B cells expressing stereotyped IGHV14-4/IGKV4-59 BCRs. Thus, K03.12 feedback did not promote the selection of entirely novel GC B cells but instead advantaged B cells normally disfavored in the competitive GC environment. While the frequency of RBS+RBS/Interface-mapped GC B cells increased from day 12 to day 16 in both control groups, that increase came at the expense of HA^-^ GC B cells rather than by reductions in the other HA^+^ epitopic categories. This observation raises the possibility that physiological selection of GC B cells with avid, epitope-specific BCRs are little affected by the expansions (or contractions?) of populations specific for other epitopic sites. If so, masking of one discrete epitope by Ab would not affect the immunodominance of the others.

Although several studies have demonstrated suppression of B-cell entry into GCs by Ab feedback, less is known about how Ab feedback affects active GC responses. Earlier work using passive IgM rAb to 4-hydroxy-3-nitrophenylacetyl at the onset of GC responses showed accelerated affinity maturation^9^. In more recent studies, the numbers of high affinity BCR KI cells in primary GCs were reduced by production of early high affinity Ab, whereas lower affinity Ab showed no effect^71,72^. In our study, substantial (41%-54%) and high affinity GC responses to stem epitopic sites would be expected to effect similar Ab feedback responses after local plasmacyte differentiation. We did not, however, observe any obvious differences in either the frequency or avidity of stem-directed GC B cells from day 12 to -16. Any early feedback effects on GC responses by endogenous Ab were too small to detect. Under physiological conditions, the quantity and avidity of early Ab may be insufficient to mediate feedback effects on primary GCs for a period determined by local Ab levels and affinity.

The distributions of AvIn values for individual epitopic categories were very broad, with ranges of three to four logs, showing primary GCs supported both high and low avidity B cells that react with the individual epitopes present on the native immunogen. As both low and high affinity B cells must compete for Tfh survival and proliferation cues^2,29^, BCR avidity for native antigen cannot be the only determinant of recruitment or persistence of GC B cells. Previous studies from our group and others showed permissive selection, the proliferation, hypermutation, and affinity maturation of B cells which are either unspecific^31,73^, have marginal avidity^74^ or are haploinsufficient in MHCII^75^. To our surprise, epitope masking by circulating Ab did not select against the entry of lower avidity B cells in the GC reaction but rather blocked GC entry or affinity maturation of higher avidity populations. This result appears counterintuitive for feedback mechanisms relying on competition between passive Ab and the available BCR repertoire^3,7,8,10^. Mutation frequencies in those low avidity cells were like those in companion GC B cells, suggesting that they had undergone authentic Tfh-mediated selection, proliferation, and mutation in GCs without improving BCR affinity to the HA immunogen.

How are low avidity B cells recruited into GC responses? A recent study noted a “complementary” role for Ab acting as an antigen scaffold to increase ligand valency and promote activation and antigen presentation by low affinity B cells^67^. Our own study describes a “cooperative” role for Ab in the formation of ICs, and creation of novel epitope structures. Those two mechanisms provide explanations for the significant increases in HA^-^, “unspecific” GC B cells present in mice receiving RBS Ab^76^. BCR repertoire differences between the “complementary” and “cooperative” mechanisms for “unspecific” B-cell recruitment are significant: low affinity B cells activated by antigen scaffolds are specific but not avid for nominal antigen whereas B cells selected on IC neoepitopes can be avid for the neoepitope but have low affinity for native antigen. Affinity-driven selection for IC neo-epitopes offers a potential explanation for the persistence and somatic evolution of GC B cells with little or no apparent affinity for the native immunogen^31^.

Given the potential for feedback activity by Ab production during primary GC responses^71,72^, generation of ICs on FDCs may diversify GC repertoires. We note that the rare IGHV14-4/IGKV4-59 GC B cells from control mice were recovered only on day 16, in the presence of endogenous H3/KS Ab^31^. The “replacement”^77^ of founder GC B cells and subsequent generation of “unspecific” memory B cells without affinity for the eliciting immunogen^78^ may be explained by recruitment of GC B cells specific for IC neoepitopes. We propose that endogenously generated ICs constitute a form of “dark antigen” that drives an increasingly diverse GC BCR repertoire over time^31^.

As musinized K03.12 contains human variable domains, it is possible that this xenogenic difference artifactually promotes the immunogenicity of K03.12 and recruitment of IGHV14-4/IGKV4-59 GC B cells. This possibility, however, is remote for at least five reasons. First, induction of Abs targeting mouse-human Ab chimeras requires repeated immunization in adjuvant^79^; our passive transfer is a single *i.v*. injection of soluble, musinized K03.12 Ab. Second, the decay rates of musinized K03.12 post-transfer were linear from day 2-31 indicating no evidence for accelerated loss due to removal by endogenous Ab specific for the K03.12 V-domains. Third, we isolated GC B cells expressing IGHV14-4/IGKV4-59 BCR from HSV Ab control groups and confirmed their structure matches that for enhanced binding to H3/KS:K03.12 complexes. Fourth, the germline L-chain residues (Y32_LC_ and S67_LC_) of IGKV4-59 that contact the H-chain V-domain K03.12 are insufficient for measurable avidity to K03.12 Ab in the absence of H3/KS. Finally, elicitation of autologous Ab responses to create endogenous ICs has been reported in animals repeatedly immunized with HIV-1 Env^80^ or a malaria vaccine^81^. Ab responses to IC are natural phenomenon and not an artifact of the chimeric musinized K03.12 Ab.

By the definitions of Brown *et al*., the IGHV14-4/IGKV4-59 Abs we have described represent class II IC Abs^80^. We think it is likely that endogenous IC Abs targeting other epitopic sites may have been induced in our experiments as we note that despite their generally low affinity for uncomplexed H3/KS, IGHV14-4/IGKV4-59 GC B cells in control group animals appeared on day 16, in the presence of endogenous serum Ab. Brown *et al*. have commented on the possibility that endogenous Ab responses to IC constitute a form of Jerne’s idiotypic network; we would emphasize that Ab feedback is not just epitopic masking but the creation of IC neoepitopes and an interacting network Ab V-domains that can modulate GC B-cell repertoires^80,82^.

This potential for modulating GC B cell repertoires may have significant effects in studies that follow GC populations over long periods or employ repeated, especially closely spaced, immunizations. In studies that follow GC populations for extended periods, a common observation is the recruitment of B cells with low or unmeasurable BCR affinities for the eliciting antigen^77,83^. This recruitment occurs in the presence of residual, high affinity GC competitors; a paradox noted by Hagglof *et al*., “*the affinity threshold for late GC entry is lowered in the presence of high-affinity antibodies*”^83^. A simple explanation for this conundrum is that the late GC entrants are not specific for the eliciting antigen *per se* but for its IC form with endogenous Ab. Repeated immunization is often employed in vaccine studies designed to elicit rare Ab specificities^80,81,84,85^. In these studies, it is not unusual for desired Ab types to appear but then be lost as serum Ab responses go “off target”. Our findings and those of others predict that as concentrations of a particular Ab type increases, *e.g*., targeted to a neutralizing epitope, *de novo* recruitment for that same epitope is diminished and resulting IC neoepitopes promote “off target” B-cell recruitment^80,81,84,85^. Vaccine strategies for epitope focusing may, by initial success, become self-defeating.

It is notable that SARS-CoV-2 boost immunization promotes expansion of memory B-cell populations specific for conserved but subdominant RBD epitopes at the cost of masking epitopes that dominated earlier responses. The helpful outcome of this result is often the elicitation of Abs with broad effect^8,22–25^. This result emphasizes the importance of balancing the potential benefits and disadvantages of Ab feedback (*i.e*., epitope masking vs IC neoepitopes) to optimize vaccine strategies. To this end, the relative locations of prime and boost immunization should be considered, as preexisting, local innate^86^ and adaptive immunity^66^, including persistent GCs, are known to aid in the recruitment or re-activation of the progeny of primary GC responses long maintained by affinity-driven selection for native Ag^37^.

## Limitations of the study

Our study sample of 33,536 clonal GC B cell cultures for repertoire analysis is significantly larger than prior studies; nonetheless, it represents only 0.3% of all GC B cells elicited in our study. This study of Ab-feedback is based on a supraphysiological amounts of passive, epitope specific IgG Ab and may not accurately describe the feedback effects of physiological concentrations of polyclonal Ab. Finally, in contrast to the fixed decay rate of passive IgG Ab, physiological Ab levels are sustained by long-lived plasmacyte populations; these longer Ab exposures were not tested in our study.

## Resource availability

### Lead contact

Requests for resources and reagents should be directed to Garnett Kelsoe.

### Materials availability

Plasmids or proteins of rAbs constructed in this study (RL1122-05606, RL1125-06507, RL1125-10403, RL1125-14202, RL1125-18706, RL1161-16405, RL1162-25608, HL1162-65809, RL1165-18604) are available upon request.

### Data and code availability

EM maps for the RL1122-05606 or HL1162-65809 immunocomplexes are deposited in the EM Data Bank (accession: EMD-77240, EMD-77237); the corresponding atomic coordinates are deposited in the Protein Data Bank (accession: 35WC, 35VS).

BCR sequences of rAbs constructed in this study (RL1122-05606, RL1125-06507, RL1125-10403, RL1125-14202, RL1125-18706, RL1161-16405, RL1162-25608, HL1162-65809, RL1165-18604) have been deposited in GenBank (accession: PZ718880-PZ718897)

## Acknowledgments

We thank our laboratory colleagues for assistance and discussion. We thank R. Abbott, K. Boradia, L. Zhu, and J. Li, for production and provision of HA molecules. We thank S. Slater and E. Kosher for technical support in flow cytometry at the Duke Human Vaccine Institute Flow Cytometry Facility (Durham, NC). Research was conducted using equipment and services of the Harvard Cryo-EM Center for Structural Biology (supported in part by the Nancy Lurie Marks Family Foundation).

Some molecular graphics were rendered with ChimeraX, developed by the Resource for Biocomputing, Visualization, and Informatics at the University of California, San Francisco, with support from NIH R01-GM129325 and the Office of Cyber Infrastructure and Computational Biology, National Institute of Allergy and Infectious Diseases.

This work was supported in part by NIH awards P01 AI089618 (SCH, GK) and K99AI193241 (JF). JF is supported by a Career Development Fellowship Award from the Basic & Clinical Translational Research Executive Committees of the Office of Faculty Development of Boston Children’s Hospital.

## Author contributions

Conceptualization, M.K., G.K.; investigation, K.T., J.F., D.L., X.L., L.Y., S.S., M.K.; data curation, K.T., E.J.S., E.V.I.; methodology, K.T., J.F., M.K., K.R.M., S.C.H., G.K.; resources, E.J.S., E.V.I., J.S.M.B., D.L., X.L., K.R.M., K.W., S.C.H.; software, E.J.S., E.V.I., J.S.M.B., K.W.; visualization, K.T., J.F., M.K., K.R.M.; writing-original draft, K.T., J.F.; writing, review, editing, J.F., E.V.I., J.S.M.B., M.K., S.C.H., G.K.; funding acquisition, J.F., S.C.H., G.K.; supervision, S.C.H., G.K.

## Declaration of interests

NB-21.2D9 feeder cell line is licensed non-exclusively to Moderna. G.K. and M.K. are inventors of the cell line.

M.K. and G.K. are coinventors on a patent (Patent No. US-12459987-B2) assigned to Albert Einstein College of Medicine and Duke University regarding the BMPC23 HSV antibody used in this study.

## Declaration of generative AI and AI-assisted technologies

The authors declare no use of generative AI or AI-assisted technologies.

## Supplemental information

Document S1. Figures S1–S10, Tables S1–S4

## STAR★Methods

### Experimental model details

#### Mice and immunizations

Female C57BL/6 mice were obtained from the Jackson Laboratory. Mice were maintained under specific pathogen–free conditions at the Duke University Animal Care Facility; seven- to 12-week-old female mice were immunized with 20μg of H3 A/Kansas/14/2017 X-327 HA trimer (H3/KS) in Alhydrogel^®^ adjuvant 2% (InvivoGen, final concentration of 1%) in the footpad of the right hind leg. Cohorts of mice were given 20μg or 200μg of rAb by intravenous (i.v.) passively transfer. Transferred rAb were injected two days before immunization and were specific either for HSV gB glycoprotein (BMPC23)^33^ or the RBS of the H3/KS (K03.12)^32^. At 12- and 16 days after immunization, popliteal lymph nodes (pLNs) of right leg were collected for the sorting and analysis of GC B cells. To calculate half-lives of passive mAbs, naïve C57BL/6 mice were i.v. injected with 20μg or 200ug of K03.12 or BMPC23 and bled from submandibular vein at 2, 17 and 31 days after the passive transfer. For negative control, naïve mice without any passive transfer were also bled. All experiments involving animals were approved by the Duke University Institutional Animal Care and Use Committee.

### Method details

#### Expression and purification of recombinant proteins

Recombinant full-length, soluble ectodomains (FLsEs) of H3/KS (WT or point mutants^39,40^) were cloned, expressed, and purified essentially as described^88–91^. In brief, codon-optimized DNA encoding an N-terminal gp67 secretion signal, the FLsE, and a C-terminal HRV3C protease cleaving site, T4 fibritin (foldon) trimerization domain and a His_6_-tag was subcloned into the pFastBac plasmid. Corresponding bacmids were used to transfect *Spodoptera frugiperda* (Sf9) cells to produce recombinant baculoviruses. *Trichoplusia ni* (Hi5) cells were infected with the recombinant baculoviruses, and 72 hours post-infection, the HA-containing supernatant was harvested and clarified by centrifugation. HA was absorbed to Co^2+^–nitrilotriacetic acid (NTA) metal affinity resin (Takara) washed extensively with buffer A (10 mM HEPES, 150 mM sodium chloride, pH 7.5), and eluted with buffer A plus 400 mM imidazole. HA was concentrated and further purified by gel filtration chromatography using a Superdex S200 column in PBS. Concentrated HAs were stored at 4°C until use. H3/KS FLSE to be used for immunization was further incubated with HRV3C protease (Thermo Fisher Scientific) overnight at 4°C to remove the foldon and His_6_-tag, then adsorbed to Co^2+^-NTA to remove the cleaved tags and the unprocessed HA. The tag-less HA was concentrated and buffer-exchanged to PBS, then stored at 4°C until use.

DNA encoding the globular, H3/KS head domain with a C-terminal HRV3C protease cleavage site and His_6_-tag was cloned into a modified pVRC8400 plasmid and transfected into Expi293F cells using polyethyleneimine. Five days post-transfection, HA-containing culture supernatant was clarified by low-speed centrifugation and purified by immobilized metal affinity chromatography (as described above). Purified HA head was buffer-exchanged to PBS, concentrated, and stored at 4°C until use. For cryo-EM, the HA head was processed with HRV3C protease (as described above), concentrated and exchanged into PBS, then stored at 4°C.

The mouse BMPC23 and human K03.12 Abs were expressed as mouse IgG1 or IgG2c rAbs as described^38,39^. In brief, DNA encoding heavy- or light chain variable domains were cloned into expression vectors containing the constant regions of mouse IgG1, IgG2c, Igκ or Igλ. Full-length IgGs were produced by transient transfection of Expi293F cells using the Expifectamine 293 transfection kit (Thermo Fisher). Five days post-transfection, supernatants were harvested, clarified by low-speed centrifugation, mixed 1:1 with Protein G binding buffer (for IgG1) or Protein A binding buffer (for IgG2c), and incubated overnight with Pierce Protein G or Protein A agarose resin (Thermo Fisher). The resin was collected in a chromatography column, washed with binding buffer, and eluted in Pierce IgG Elution Buffer (Thermo Fisher) followed by the immediate neutralization by 1M Tris (pH 9). After the dialysis into PBS, IgG concentrations were determined with a NanoDrop spectrophotometer (Thermo Fisher).

Recombinant Fabs were expressed with a C-terminal HRV3C protease cleavage site and His_6_-tag on the heavy chain. Expression plasmids encoding the HC and LC were co-transfected into Expi293F cells using polyethyleneimine. Five days post-transfection, Fab-containing culture supernatant was clarified by low-speed centrifugation. Fab was purified by adsorption to Co^2+^-NTA resin, as described above for HAs. Purified Fab was concentrated and buffer-exchanged into buffer A. The His_6_-tag was removed by incubating Fab with His_6_-tagged HRV3C protease (ThermoFisher) overnight at 4°C, followed by adsorption to Co^2+^-NTA as described above. The flow-through fraction containing tag-less Fab was concentrated and exchanged into PBS plus 0.1% (w/v) sodium azide, then stored at 4°C.

#### Biolayer interferometry (BLI)

BLI experiments were performed on a BLItz label-free protein analysis system (ForteBIO). All measurements were in PBS at room temperature. Purified, 8xHis-tagged H3/KS head (2.6 μM) was immobilized on Ni-NTA biosensors (Sartorius), and Fab was titrated to determine association and dissociation rates. To determine binding kinetics against a complex of HA+K03.12 Fab, 8xHis-tagged H3/KS head (2.6 μM) was mixed with K03.12 Fab (4 μM, no His-tag) and incubated at room temperature for ≥10 min before being immobilized on Ni-NTA biosensors. RL1122-05606 Fab was then titrated to determine association and dissociation rates, with wash and dissociation steps carried out in the presence of K03.12 (4 μM).

#### Flow cytometry

Single-cell suspensions from recovered pLNs were dispersed by gentle disruptions between glass slides and suspended in 10% DMEM; DMEM supplemented with 10% FCS, 55µM 2-mercaptoethanol, 100 units/ml penicillin, 100μg/ml streptomycin and additional 2mM L-glutamine (all Invitrogen). Suspended cells were then pre-treated with the mixture of anti-CD16/CD32 Ab (2.4G2, BD) and rat IgG (Invitrogen) on ice to reduce unspecific interactions between cells and labeling Abs. After 30 minutes, cells were labeled with fluorophore-conjugated Abs^37^: anti-TCR β-chain BV711 (H57-597, BioLegend), anti-B220 BV785 (RA3-6B2, BioLegend), anti-CD38 PE-Cy7 (90, BioLegend), GL7 FITC (BD), anti-CD138 BV605 (281-2, BioLegend), and anti-IgD BV421 (11-26c.2a, BioLegend) for an additional 30 minutes. Stained cells were washed twice with 10% DMEM and suspended in the same media containing propidium iodide (PI, Sigma-Aldrich). GC B cells were identified as the TCRβ^-^B220^hi^CD138^-^ CD38^lo^GL7^+^ population; cell doublets were excluded by by FSC-A/FSC-H gating and dead cells by PI signal. Single GC B cells were sorted into each well of 96-well plates using BD FACSymphony S6 sorter. Flow cytometry data were analyzed using FlowJo software (Treestar Inc.).

#### Single B-cell Nojima culture

Individual GC B cells were expanded by culture on monolayers of NB-21.2D9 feeder cells as described^31,37,39^. In brief, NB-21.2D9 cells were pre-seeded (2,000 cells per well) onto 96-well plates in 100μl of B cell media (BCM): RPMI-1640 (Invitrogen) with 10% FCS (HyClone, Cytiva), 55µM 2-mercaptoethanol, 10mM HEPES, 1 mM sodium pyruvate, 100 units/ml penicillin, 100μg/ml streptomycin, and MEM non-essential amino acids (Invitrogen). On the following day, 100μl of BCM containing 2ng/ml recombinant mouse IL-4 (PeproTech) was added to every well, and shortly after, single GC B cells were sorted into individual wells. After two days of culture, media were replaced by removing 100μl of supernatant and adding 200μl of fresh BCM. From days 4 to 8, media exchange by removing and replacing 200 μl of culture media daily. On day 10, culture supernatants were harvested for ELISA and subsequent screening in multiplex bead assay. Culture plates containing GC B cell clones were stored at −80°C until use for V(D)J amplification.

#### ELISA

IgG secretion into culture supernatants was determined by ELISA^31,37,39^. In brief, 2µg/ml of anti-mouse Igκ plus -Igλ capture Abs (Southern Biotech) were coated on 384-well microplates in 0.1 M sodium carbonate buffer (pH 9), followed by blocking with PBS containing 0.5% BSA. For ELISA, culture supernatants were diluted 1:10 with PBS containing 0.5% BSA and 0.1% Tween20. Diluted supernatants (50μl) were introduced into blocked ELISA plates overnight (4° C) and then washed extensively. After washing, HRP-conjugated goat anti-mouse IgG secondary Ab (1:5000 in PBS with 0.5% BSA and 0,1% Tween 20; Southern Biotech) was added and bound HRP activity visualized using a TMB substrate kit (BioLegend). OD values at 450nm were read by Spectramax plate reader (Molecular Devices).

Binding of recombinant mAbs against K03.12 or BMPC23 (both human IgG1) was assessed by coating 2µg/ml human mAbs and serially diluting mouse IgG samples (starting 1μg/ml; seven points of 3-fold dilutions) in PBS containing 1% BSA, 0.05% NaN_3_, 0.05% Tween 20 and 1% non-fat milk, followed by the same protocol as above. Mouse anti-human IgG (Fcγ fragment specific, Jackson ImmunoResearch Inc.) capture Ab was used as a positive control.

#### Beads-based multiplex assay (Luminex assay)

Antigen specificity and avidity of IgG in culture supernatants was determined in a Luminex assay as described^31–33,37–39,51,92^. In brief, culture supernatants were diluted 1:10 in PBS containing 1% BSA, 0.05% NaN_3_, 0.05% Tween 20 and 1% non-fat milk and incubated with the mixture of antigen-coupled microsphere beads in 96-well filter bottom plates (Millipore) overnight. After washing, beads were incubated with 2μg/ml of PE-conjugated goat anti-mouse IgG (Southern Biotech) for one hour and analyzed on a Bio-Plex 3D Suspension Array System (Bio-Rad). The antigens were coupled to carboxylated Luminex beads (Luminex Corp): BSA, goat-anti mouse IgG, goat anti-mouse Igκ, goat anti-mouse Igλ (all Southern Biotech), and rHAs (both WT and point mutants).

Antigen specific binding of serum Abs or recombinant mAbs was assessed by serially diluting samples (starting 10- to 100-fold and 2μg/ml, respectively; seven points of 4-fold dilutions) in PBS containing 1% BSA, 0.05% NaN_3_, 0.05% Tween 20 and 1% non-fat milk, followed by the same protocol as above.

For competition assays, beads were incubated with human IgG1 version of K03.12 or S5V2-29 with dilution (starting 10μg/ml; 11 points of 3-fold dilutions) in PBS containing 1% BSA, 0.05% NaN_3_, 0.05% Tween 20 and 1% non-fat milk overnight. For no competitor control, BMPC23 was utilized in the same manner. Then, without washing, mouse IgG1 version of sample mAbs were added to beads at the defined concentrations in the same buffer (RL1122-05606, 300ng/ml; RL1125-06507, 1μg/ml; RL1125-10403, 20ng/ml; RL1125-14202, 300ng/ml; RL1125-18706, 300ng/ml; RL1161-16405, 100ng/ml; RL1162-25608, 10ng/ml; HL1162-65809, 20ng/ml; RL1165-18604, 1μg/ml; K03.12, 20ng/ml; S5V2-29, 1μg/ml) and incubated for two hours. After washing, beads were incubated with 2μg/ml of PE-conjugated rat anti-mouse IgG1 (BioLegend) for one hour and analyzed as above.

#### Amplification and analysis of V(D)J rearrangements

Clonal V(D)J rearrangements recovered from GC B-cell cultures were amplified and sequenced as described^37^. In brief, total RNA was extracted from selected samples using the Quick-RNA 96 Kit (Zymo Research) and subsequently treated with deoxyribonuclease I. cDNA was synthesized from recovered RNA using SMARTScribe Reverse Transcriptase (Clontech) with 0.2μM each of gene-specific reverse primers and 1μM of 5′ SMART template-switching oligo that contained plate-associated barcodes. cDNA was then subjected to two rounds of semi-nested PCRs using Herculase II fusion DNA polymerase (Agilent Technologies) with combinations of forward primers and reverse primers containing well-specific barcodes. The V(D)J amplicons were gel-purified, pooled and submitted to DNA Link Inc. or CD Genomics Inc. for DNA sequencing using the PacBio SMRT or Nanopore sequencing platform. Individual reads were processed using the same pipeline previously described for analysis of Pac Bio reads^93^. Immunogenetic annotation of the final refined consensus sequences and downstream filtering was performed with a custom pipeline built on top of Partis^94^ which was previously described^95^. Initial heavy chain clonal inference was performed as described previously, but integration of the light chain information with heavy chain clones was done as follows. Secondary clone identifiers were assigned based on light-chain features. Since somatic hypermutation can introduce sequence variation among clonally related light chains, exact light-chain CDR3 sequence matching was not required. Instead, sequences within the same heavy-chain clone were assigned to the same secondary clone when they shared the same light-chain V gene, J gene, and CDR3 length. This approach provided a conservative feature-based strategy for identifying light-chain-associated substructure while avoiding over-fragmentation of clonally related sequences due to SHM-associated sequence differences. If more than one secondary clone identifier was associated with a heavy chain clone, the features of the most frequently observed secondary clone were used as the representative light chain features for that clone. Finally, the obtained heavy and light chain paired sequences were determined for their V(D)J rearrangements and mutations using IMGT/V-QUEST (www.imgt.org/), followed by clonal assignments using Cloanalyst^52^.

#### Cryo-EM sample preparation

H3/KS head was incubated with 2-fold molar excess of RL1122-05606 Fab, 1.2-fold molar excess of K03.12 Fab, 1.2-fold molar excess of S8V1-157 Fab and 0.1% (w/v) octyl-β-glucoside (Thermo Fisher) in PBS at 4°C for ≥30 min. The total protein concentration was 2.8 mg/ml. 3.5 µL of sample was deposited onto 300 mesh Quantifoil Cu 1.2/1.3 grids that had been glow discharged in a PELCO easiGLOW (Ted Pella) at 0.39 mBar, 15 mA for 30 s. Samples were vitrified in 100% liquid ethane using a Vitrobot Mark IV (Thermo Fisher Scientific), with a wait time of 2 s, blot time of 11 s and a blot force of 5 at 22°C and 100% humidity.

The sample comprising HL1162-65809 Fab plus H3/KS head, K03.12 Fab, and S8V1-157 Fab was prepared similarly, except that HL1162-65809 Fab was supplied at 1.5-fold molar excess compared to HA head, and the total protein concentration was 2.5 mg/ml.

#### Cryo-EM data collection and processing

Cryo-EM data were recorded on a 300 kV Titan Krios G3i Microscope (Thermo Fisher Scientific) equipped with a Falcon 4i detector and a Selectris energy filter (10 eV slit width) at the Harvard Cryo-Electron Microscopy Center for Structural Biology at Harvard Medical School. Data were acquired in counting mode using the automated data collection software EPU (Thermo Fisher Scientific). Details of the data collection and dataset parameters are summarized in Table S4.

All data processing was performed using CryoSPARC^96^. Dose-fractionated images were gain-normalized, aligned, dose-weighted and summed, and then patch contrast transfer function (CTF) and defocus value estimation were performed.

For the RL1122-05606 immunocomplex, the CryoSPARC “blob” picker identified 4,684,294 putative particles, of which a random subset comprising ∼220,000 particles was used for ab-initio reconstruction of four classes, producing three junk classes and one class showing density clearly corresponding to three Fabs bound to the HA head. Junk particles were then culled from the complete dataset by several rounds of heterogeneous refinement using the output classes from ab-initio reconstruction. Selected particles were used to train a Topaz particle-picking model^97^, which then picked new particles from all micrographs in the dataset. These particles were pooled with those identified by the blob picker, duplicate particles were removed, and additional rounds of heterogenous refinement were used to cull junk particles. The final dataset, comprising 587,026 particles, was subjected to non-uniform refinement^98^ with reference-based motion correction^99^ to produce the final ∼2.9Å EM map.

A similar strategy was used to process the cryo-EM data for the HL1162-65089 immunocomplex. The final dataset, comprising 256,207 particles, was subjected to non-uniform refinement with reference-based motion correction to produce the final ∼3.2Å EM map.

#### Atomic model building and refinement

ModelAngelo^100^ was used to build atomic models of the Fabs and HA from the cryo-EM map. Parts of the atomic models that were not built by ModelAngelo – but for which there was sufficient map density to justify placement of additional residues – were built manually in Coot^101^ (v0.9.8.91). The constant domains of the Fabs were ignored, due to low density at these regions of the map. The atomic model was refined with the ISOLDE plugin^102^ of UCSF ChimeraX^103^ (v1.9) and with PHENIX^104,105^ (v1.21.1-5286). Martin numbering was applied to the Fab residues with the Abnum server^106^. PyMOL^107^ (v3.1.8) was used for visualization and analysis of the final model. Structural biology applications used in this project were compiled and configured by SBGrid^108^.

#### Statistics

Statistical significance (*P* < 0.05) was determined by Kruskal-Wallis test with Dunn’s multiple comparisons using GraphPad Prism software (GraphPad Software). Statistic testing is indicated within each figure legend.

## Supplementary figure titles and legends

**Figure S1.**
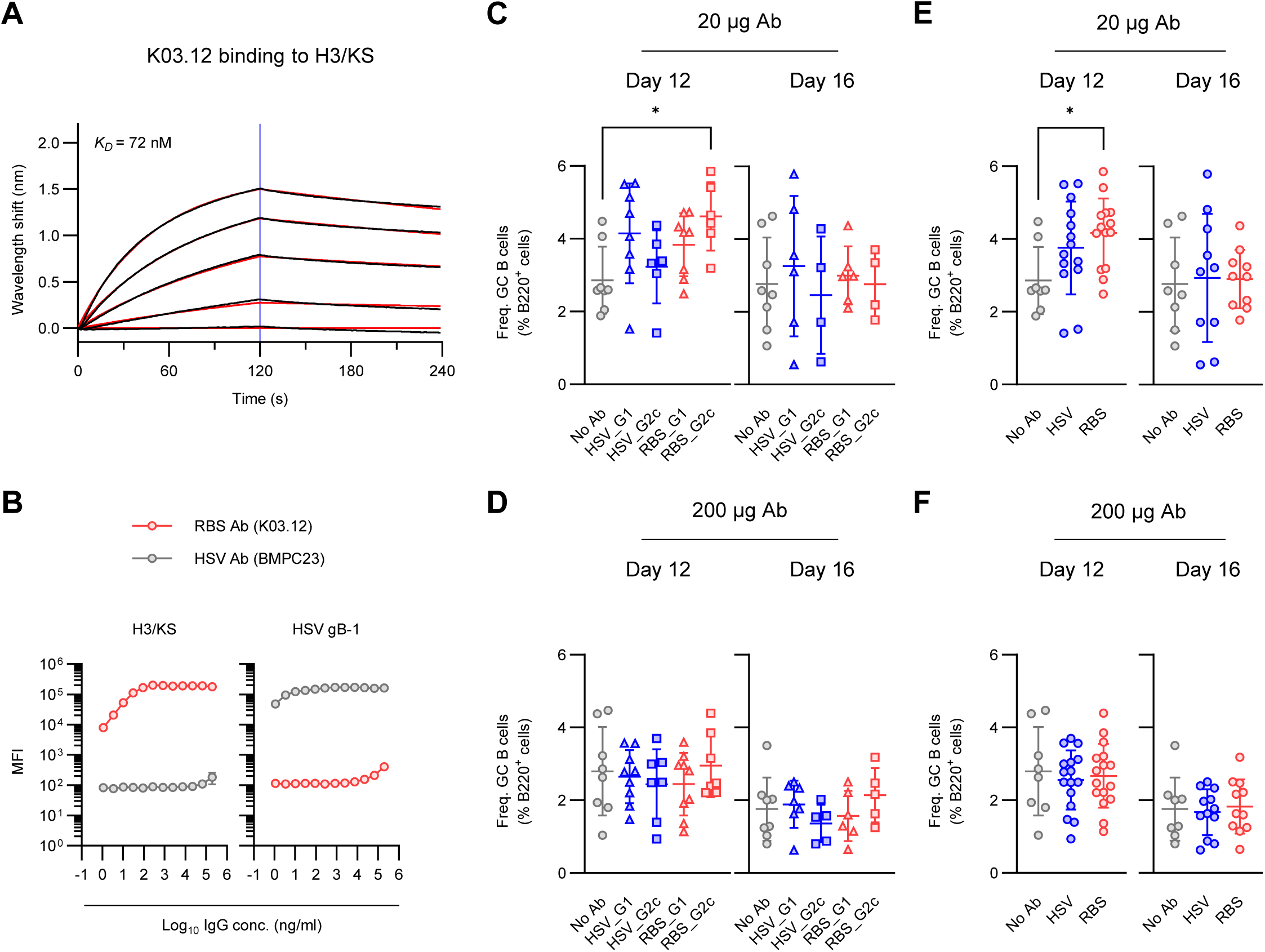
Evaluation of GC size under the passive Ab transfer, related to Figure 1. (A) Affinity measurement of K03.12 against H3/KS. Kinetics of binding intensity was measured using BLI. Measured and fitted curves are shown in black and red, respectively. Blue line segregates the association (left side) and dissociation (right side) curves. Calculated K_D_ value is shown at left top of the graph. (B) Binding ability of K03.12 (red) and BMPC23 (gray) against H3/KS or HSV gB-1 protein at high concentrations. MFI values measured by Luminex assay are plotted. Binding of each mAb was measured in a duplicate manner. Data are shown by mean±SD. (C-F) Calculated frequencies of GC B cells among total B cells are shown by (C and D) separating groups transferred with IgG1 (triangle) and IgG2c (square) or (E and F) pooling those IgG subclasses (circle). Each plot represents individual animals from No Ab (gray), HSV Ab (blue), or RBS Ab (red) cohorts with (C and E) 20µg or (D and F) 200µg of passive transfer. The data are pooled from 4-8 experiments (n=4-8 biological replicates without combining IgG subclasses). Bars indicate mean±SD. *P*-values were calculated by Kruskal-Wallis test with Dunn’s multiple comparisons: \**p* < 0.05.

**Figure S2.**
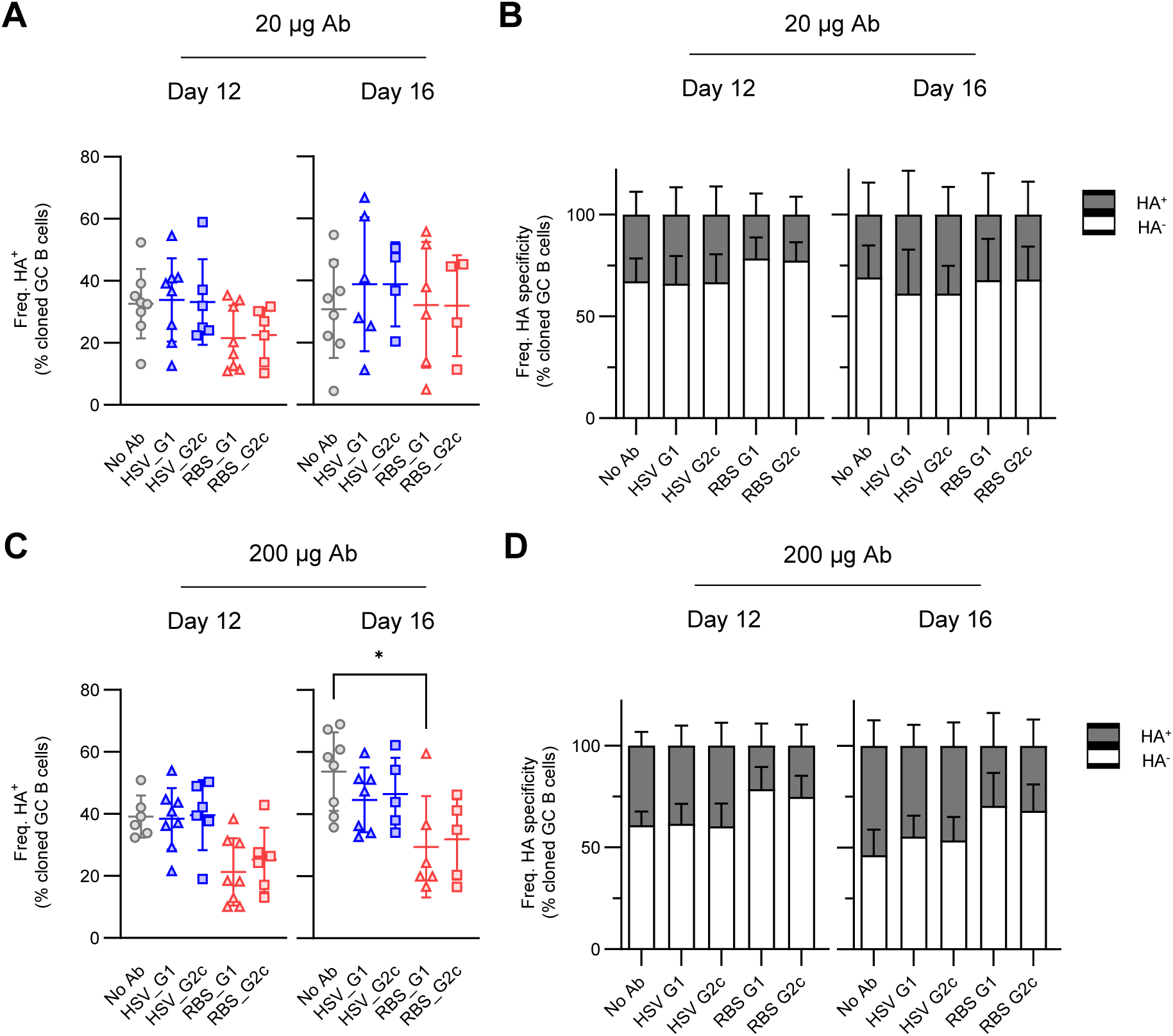
Frequency of HA-specific GC B cells shown separately by IgG Fc-subclasses, related to Figure 2. (A and C) The frequency of H3/KS reactive GC B cell cultures among total IgG secreting clonal cultures. Each plot represents individual animals from No Ab (gray circle), HSV Ab IgG1 (blue triangle) or -IgG2c (blue square), or RBS Ab IgG1 (red triangle) or -IgG2c (red square) cohorts with (A) 20µg or (C) 200µg of passive transfer. Bars indicate mean±SD. *P*-values were calculated by Kruskal-Wallis test with Dunn’s multiple comparisons: \**p* < 0.05. (B and D) The ratio of H3KS specific (gray) and -unspecific (white) GC B cell cultures among total IgG secreting clonal cultures. Bars indicate mean+SD of HA^+^ or HA^-^ population in each group. (A-D) The data are pooled from 4-8 experiments (n=4-8).

**Figure S3.**
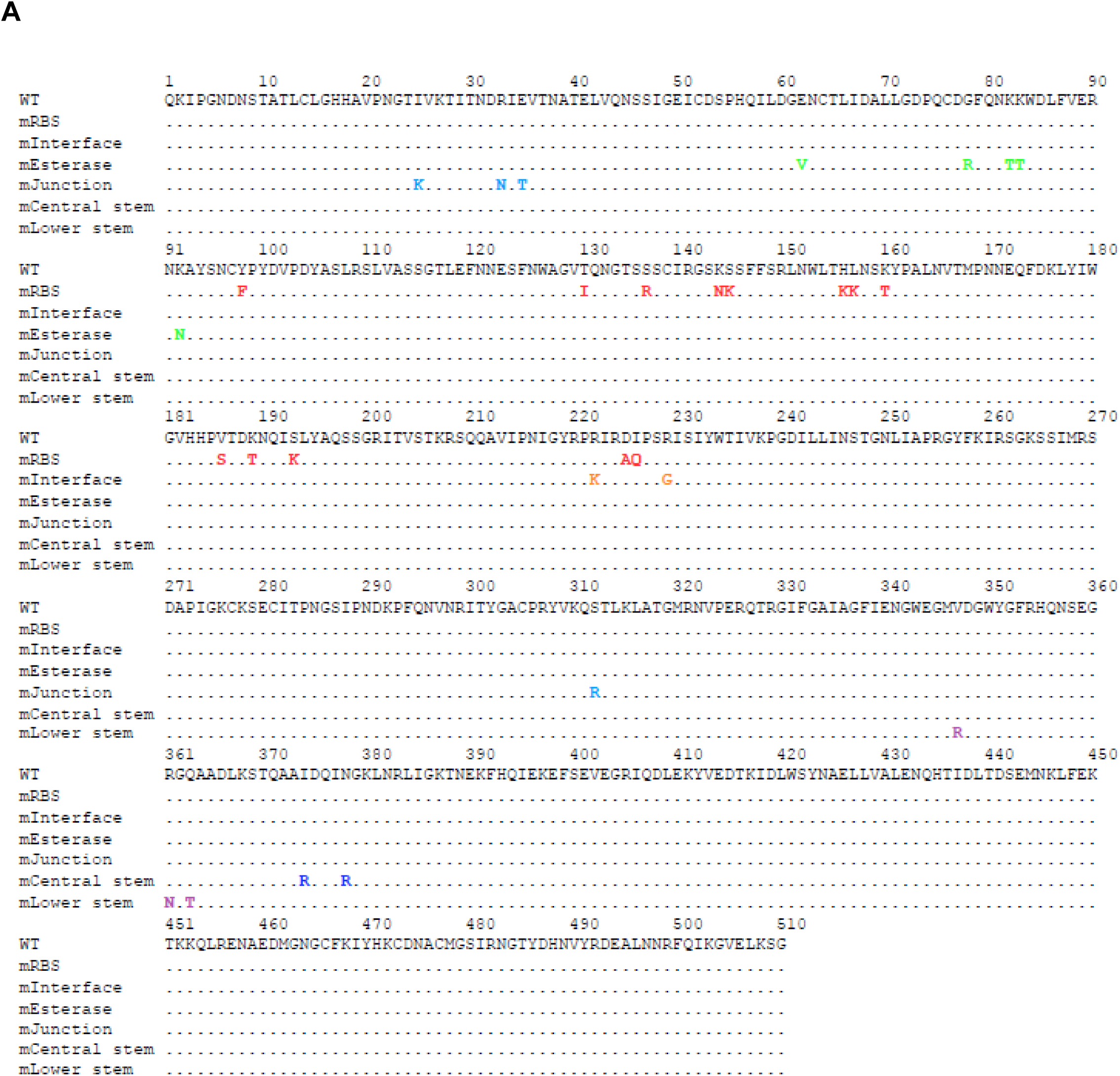
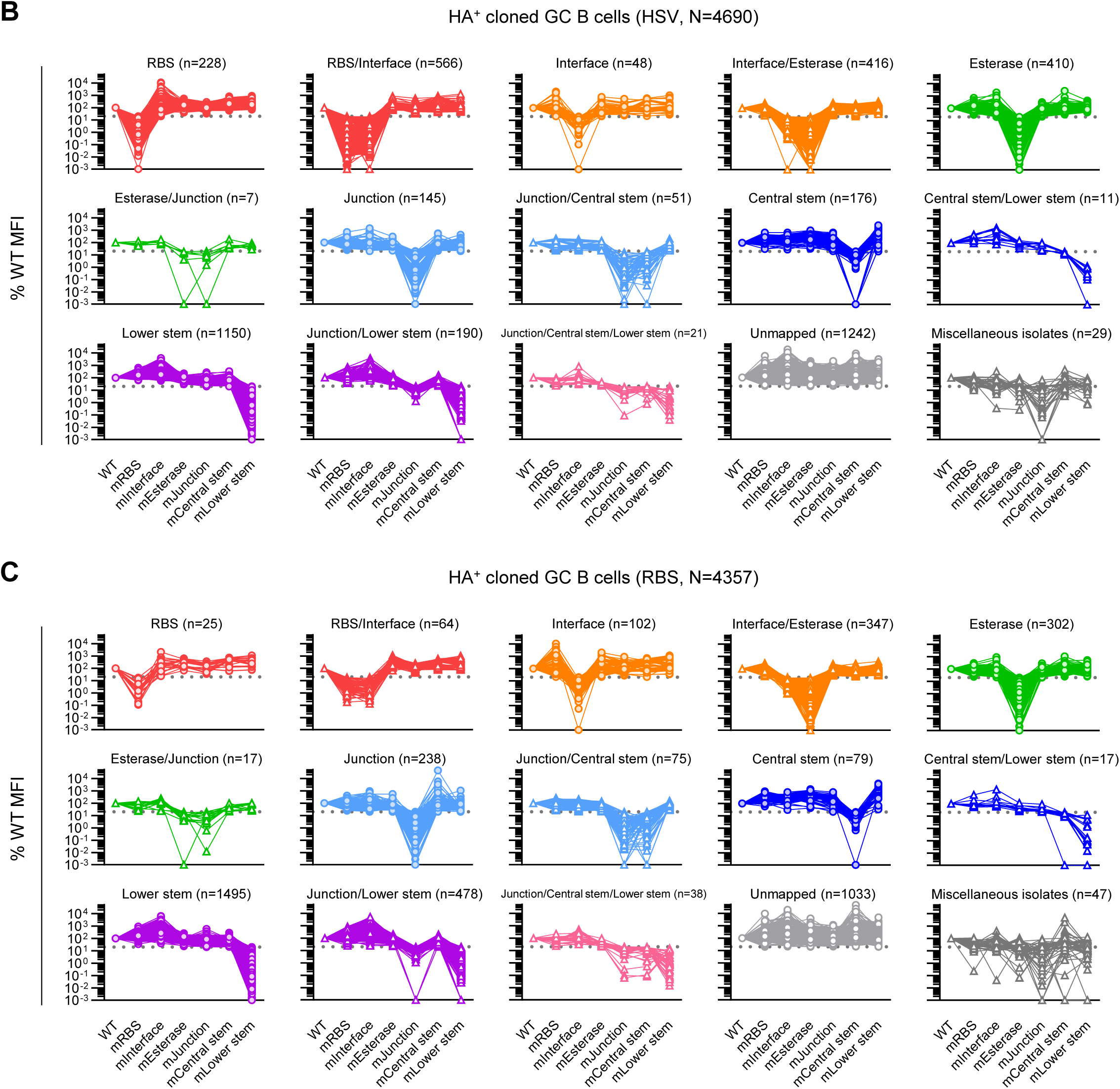
Identification of 15 epitopic categories in GC B cells induced by H3/KS under passive Ab transfer, related to Figure 3. (A) Sequence alignment of H3/KS constructs. Amino acid sequence of WT H3/KS ectodomain is shown on the top of each alignment and numbered by H3 numbering. For mutants, only mutated residues are shown in different colors each of which is matched with Figure 3A. (B and C) Evaluation of the H3/KS mutant panel using HA^+^ GC B cell cultures. The percentages of MFI values relative to the WT H3/KS construct are shown. Each plot represents individual cultures, and values from the same sample are connected with a line. HA specific cultures of (B) HSV Ab and (C) RBS Ab group obtained in Figure 2 (N=4,690 and 4,357, respectively; pooling both low and high dose conditions from day 12 and day 16 samples) were divided into 15 epitopic categories, and each category of GC B cell cultures is shown separately with the number of corresponding cultures on the top of individual figures. Specific reduction was set by 80% reduction and the threshold line (20% of WT MFI) is indicated by dotted lines. The cultures which were mapped by two or more mutants but not assigned by any of 13 mapped categories were pooled and defined as miscellaneous isolates (dark gray).

**Figure S4.**
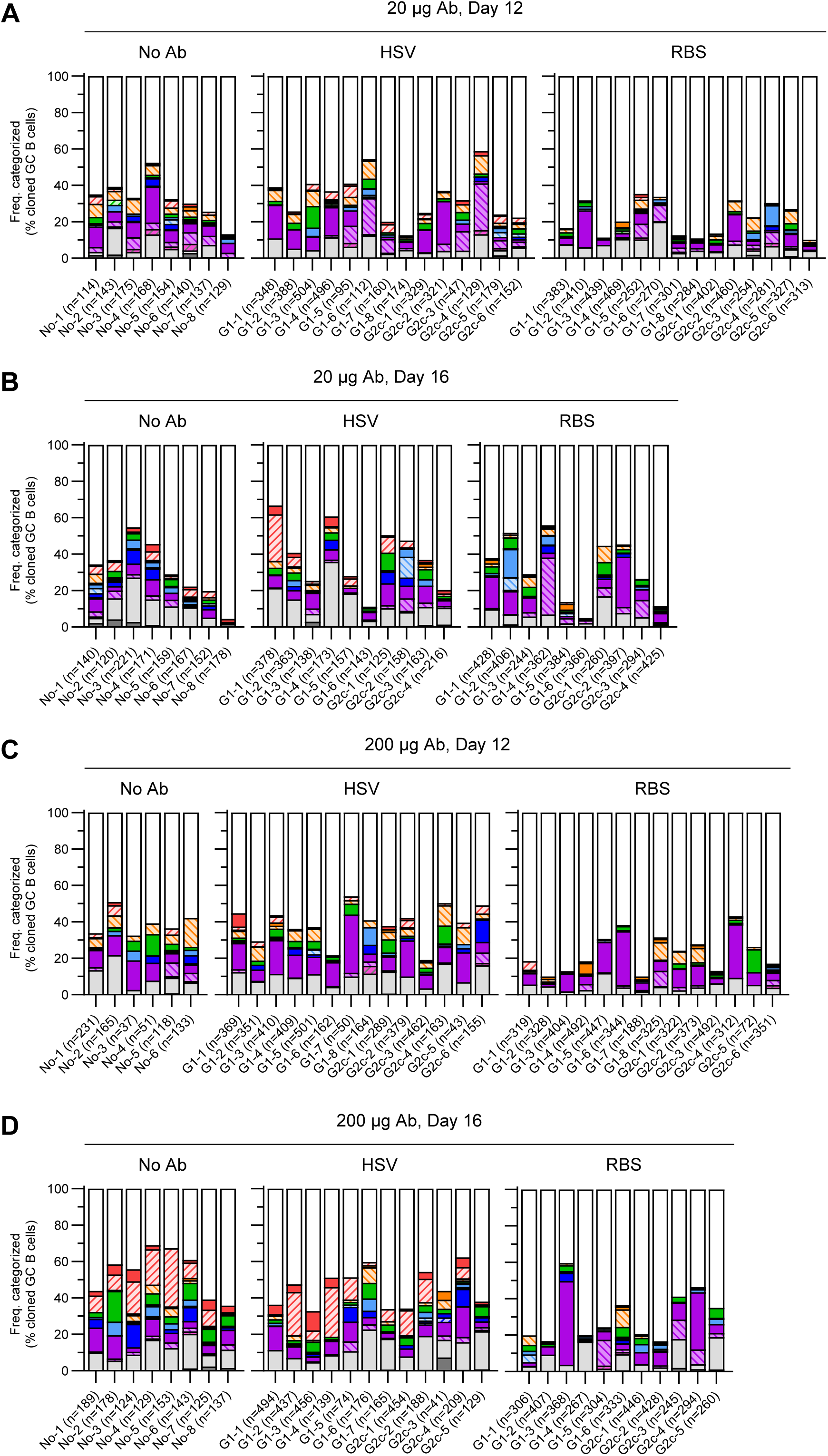
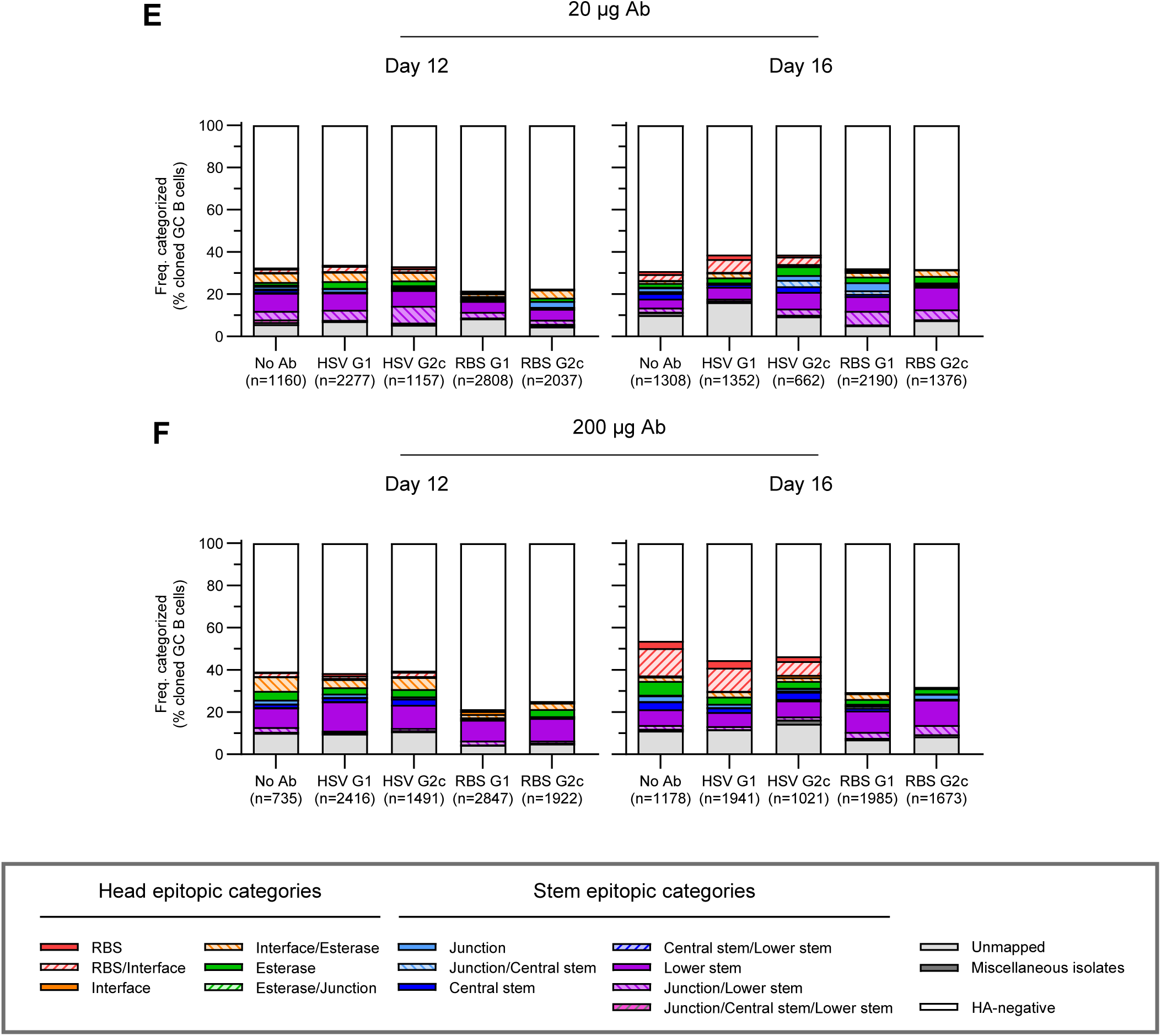
Distribution of epitopic categories in GC B cell cultures, related to Figure 4. (A-F) Frequency distribution of 15 epitopic categories among total IgG^+^ GC B cell cultures. HA specific clonal cultures obtained in Figure 2 were examined for their epitope specificities. The numbers of IgG^+^ cultures, including HA^-^ cultures, are indicated at the bottom of each column. Data are shown either by individual mouse (A-D) or by pooling mice in each group (E and F). (A-D) The mice given IgG1 or IgG2c subtype is indicated as G1 and G2c, respectively, at the bottom of each column. (E and F) No Ab, HSV Ab, or RBS Ab group with (E) 20µg or (F) 200µg of passive transfer are shown with separating IgG1 and IgG2c subgroups. The frequencies were calculated by individual animals (n=4-8 biological replicates) and subdivided bars represent the mean of them.

**Figure S5.**
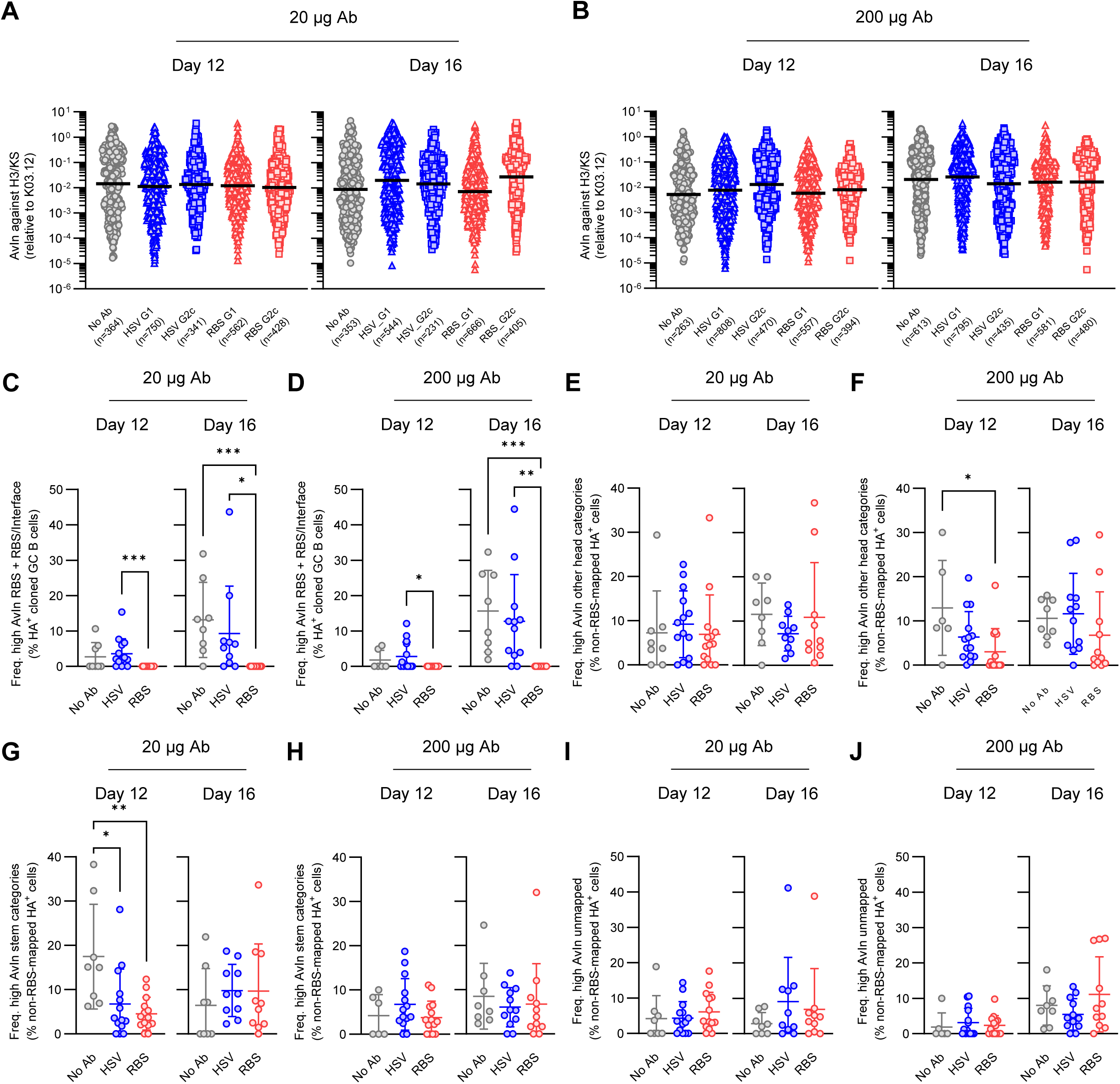
Analysis of affinity maturation in GC B cells under passive Ab transfer, related to Figure 5. (A and B) Distribution of avidities in H3/KS specific GC B cell cultures. HA specific clonal cultures obtained in Figure 2 were analyzed. The numbers of samples which were able to calculate AvIn are indicated at the bottom of each column. Each plot represents individual GC B cells from No Ab (gray), HSV Ab IgG1 (blue triangle) or -IgG2c (blue square), or RBS Ab IgG1 (red triangle) or -IgG2c (red square) cohorts with (A) 20µg or (B) 200µg of passive transfer. Bars represent median. (C and D) Frequency of high avidity GC B cell cultures mapped as RBS and RBS/Interface among H3/KS reactive population. (E-J) Frequency of high avidity GC B cell cultures mapped as (E and F) other head, or (G and H) stem, or defined as (I and J) unmapped. The frequencies were calculated with excluding cultures mapped as RBS+RBS/Interface from total H3/KS reactive population for normalization. (C-J) Each plot represents individual animals from No Ab (gray), HSV Ab (blue), or RBS Ab (red) cohorts with (C, E, G and I) 20µg or (D, F, H and J) 200µg of passive transfer (n=6-14 biological replicates with combining IgG subclasses). Bars indicate mean±SD. *P*-values were calculated by Kruskal-Wallis test with Dunn’s multiple comparisons: \**p* < 0.05, \*\**p* < 0.01, \*\*\**p* < 0.001.

**Figure S6.**
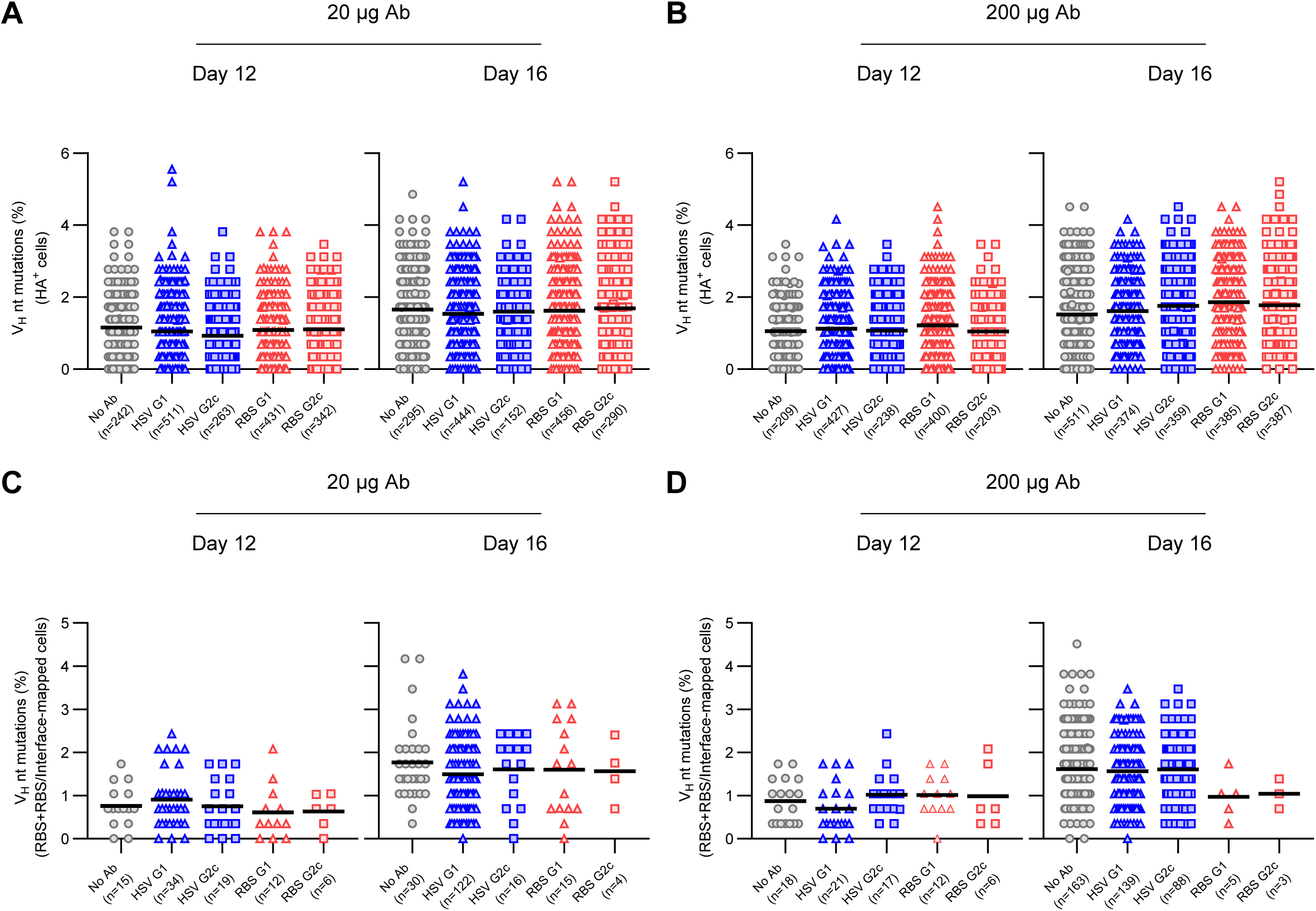
Sequence analysis of HA-specific GC B cells with separating IgG subclasses, related to Figure 6. (A and B) Distribution of V_H_ mutation frequency in H3/KS reactive GC B cell cultures. HA specific clonal cultures obtained in Figure 2 were sequenced. The numbers of samples which were successfully recovered with complete heavy plus light chain pairs are indicated at the bottom of each column. Each plot represents individual sequenced GC B cells from No Ab (gray), HSV Ab IgG1 (blue triangle) or -IgG2c (blue square), or RBS Ab IgG1 (red triangle) or - IgG2c (red square) cohorts with (A) 20µg or (B) 200µg of passive transfer. Bars represent mean. (C and D) Among the sequences obtained in (A) and (B), distributions of V_H_ mutation frequency in GC B cell cultures mapped as RBS and RBS/Interface are shown. The numbers of corresponding cells are indicated at the bottom of each column. Each plot represents individual sequenced GC B cells from No Ab (gray), HSV Ab IgG1 (blue triangle) or -IgG2c (blue square), or RBS Ab IgG1 (red triangle) or -IgG2c (red square) cohorts with (C) 20µg or (D) 200µg of passive transfer. Bars represent mean.

**Figure S7.**
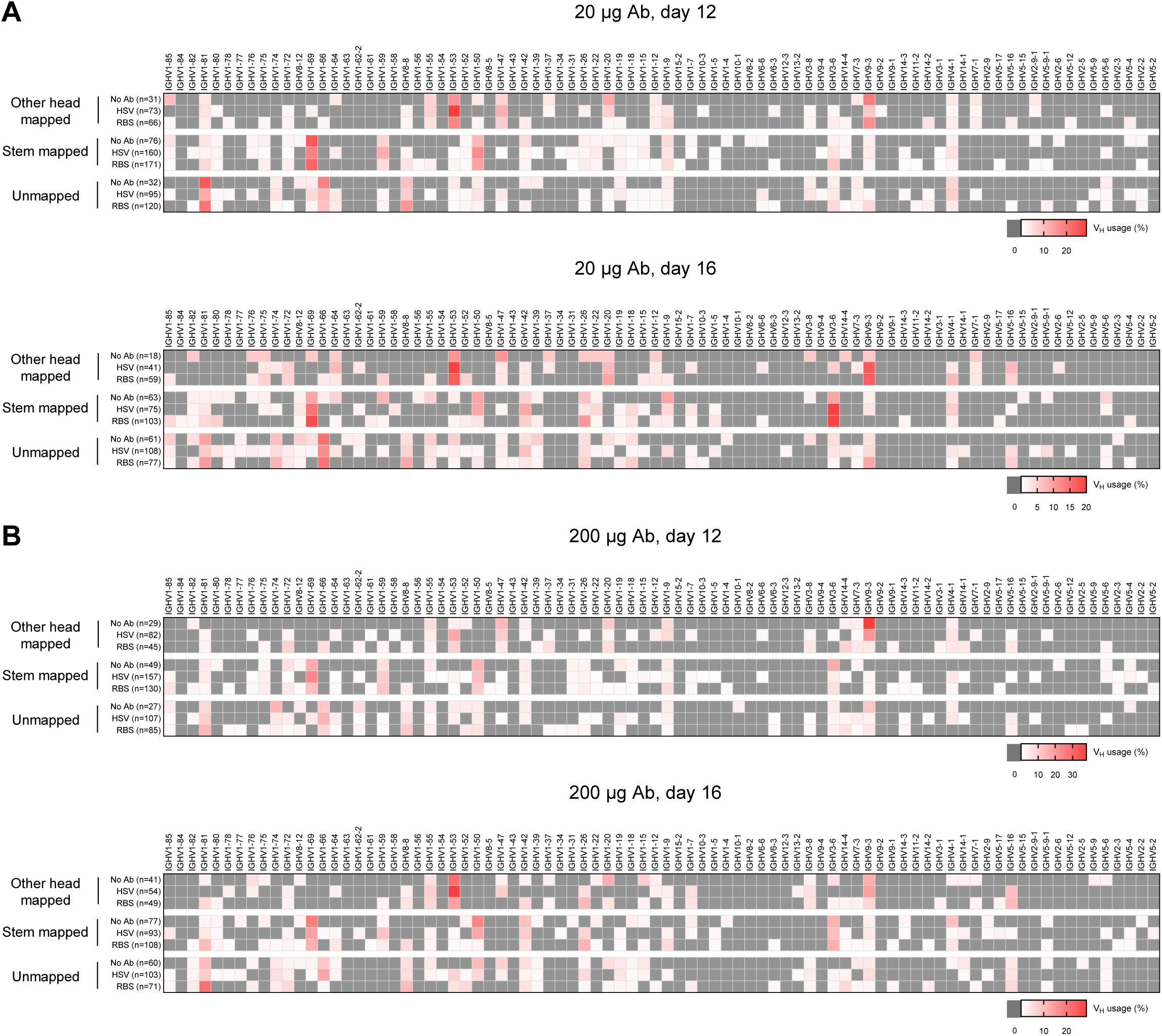
V_H_ gene usage of non-RBS-mapped GC B cells, related to Figure 7. (A and B) V_H_ gene usage of cultured GC B cells mapped as other head or stem, or defined as unmapped. Heatmaps for No Ab, HSV Ab, or RBS Ab cohorts with (A) 20µg or (B) 200µg of passive transfer are shown. The numbers of unique clones mapped as each epitopic group are indicated next to the names of groups, and the frequencies of each V_H_ gene usage were calculated by that number. Values are color-coded with higher frequency in thicker red (0% in dark gray). All the V_H_ genes identified in other head mapped, stem mapped, or unmapped GC B cells throughout the experiments are shown in the order encoded in the gene locus (left, distal to constant region; right, proximal to constant region).

**Figure S8.**
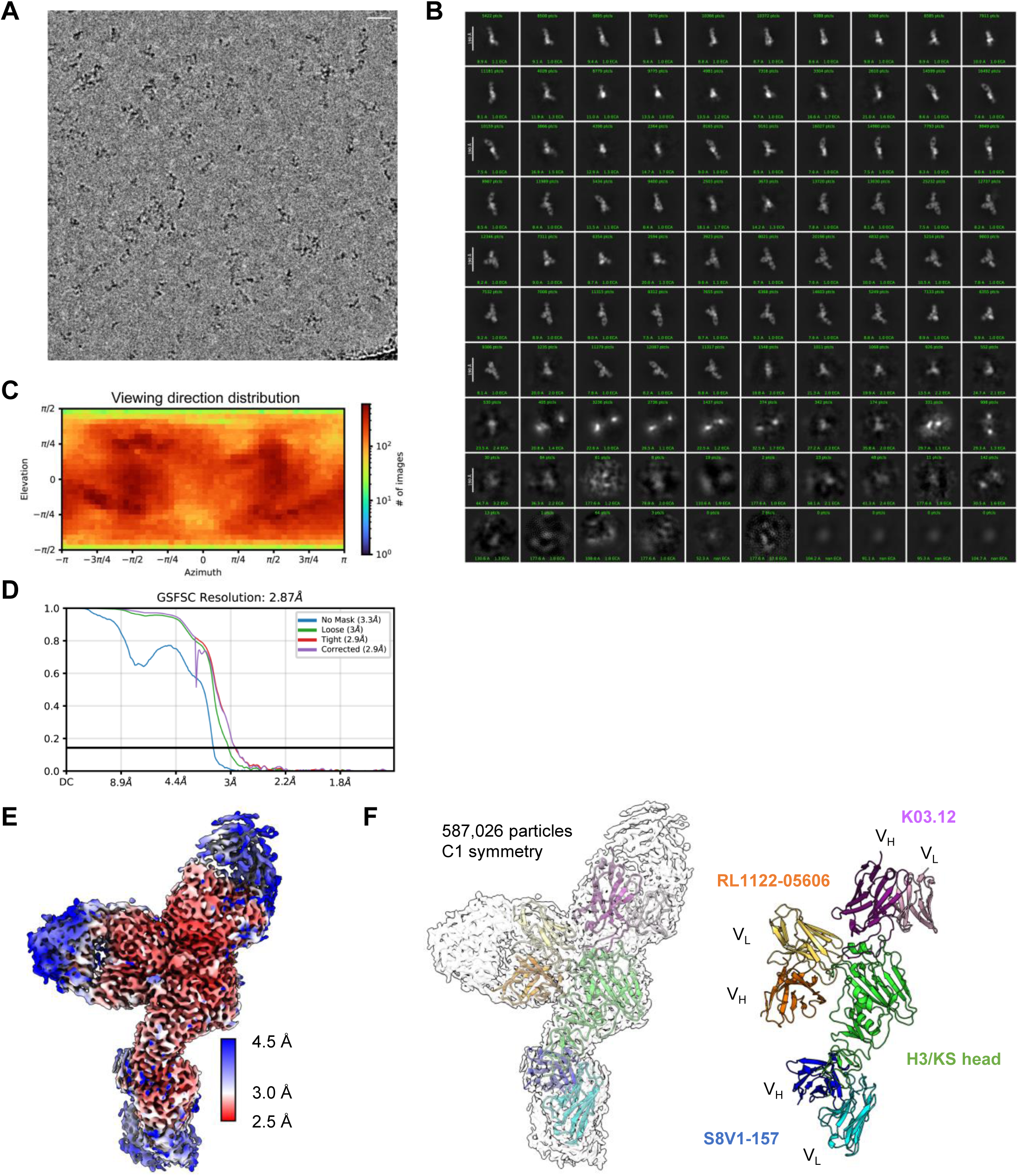
Cryo-EM analysis of RL1122-05606 Fab bound to a complex of H3/KS head plus K03.12 Fab and S8V1-157 Fab, related to Figure 8. (A) Representative electron micrograph of vitrified Fab:HA complexes, low-pass filtered with a 5 Å cutoff. Scale bar = 200 Å. (B) 2D class-averages for the particles included in the final cryo-EM map. (C) Heatmap showing the distribution of viewing directions for the Fab:HA complex. (D) Gold-Standard Fourier Shell Correlation (GSFSC) curve for the final cryo-EM map. An FSC cutoff of 0.143 was used to calculate the global resolution. (E) The final cryo-EM map, colored by estimated local resolution. (F) Left: Cartoon model of the Fab:HA complex docked into the final cryo-EM map. Right: The same cartoon model, without the cryo-EM map.

**Figure S9.**
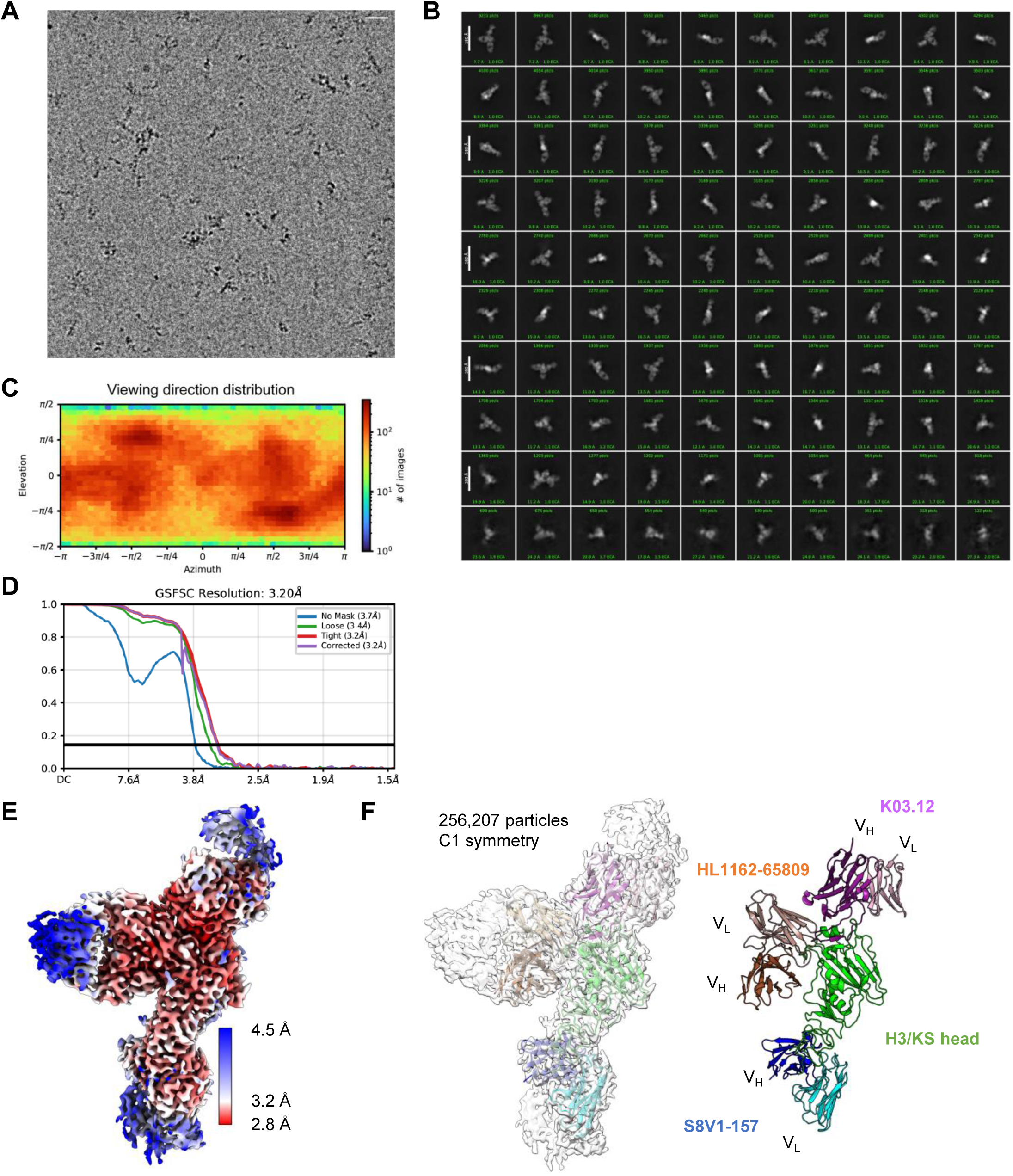
Cryo-EM analysis of HL1162-65809 Fab bound to a complex of H3/KS head plus K03.12 Fab and S8V1-157 Fab, related to Figure 8. (A) Representative electron micrograph of vitrified Fab:HA complexes, low-pass filtered with a 5 Å cutoff. Scale bar = 200 Å. (B) 2D class-averages for the particles included in the final cryo-EM map. (C) Heatmap showing the distribution of viewing directions for the Fab:HA complex. (D) Gold-Standard Fourier Shell Correlation (GSFSC) curve for the final cryo-EM map. An FSC cutoff of 0.143 was used to calculate the global resolution. (E) The final cryo-EM map, colored by estimated local resolution. (F) Left: Cartoon model of the Fab:HA complex docked into the final cryo-EM map. Right: The same cartoon model, without the cryo-EM map.

**Figure S10.**
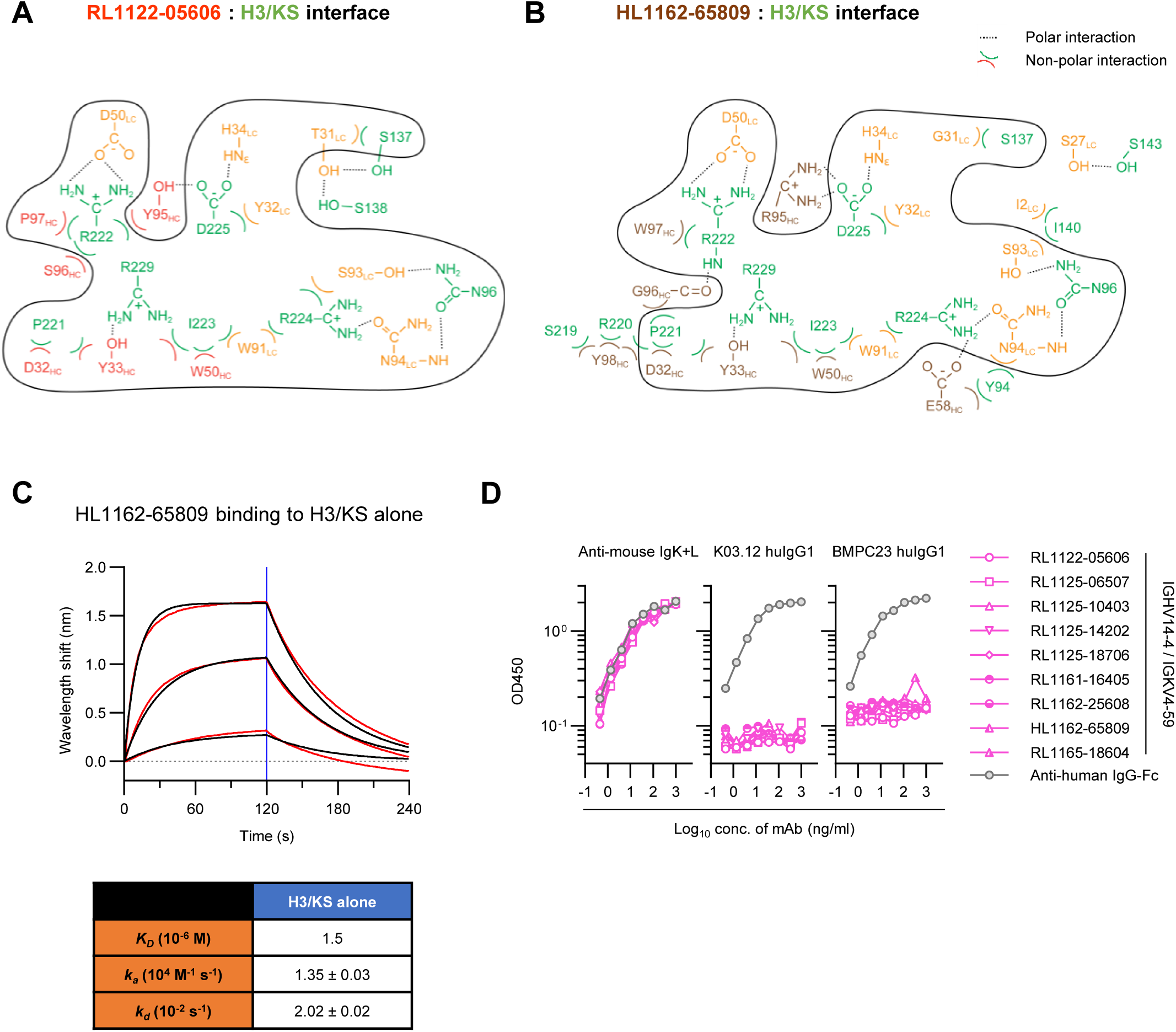
Interactions between H3/KS and RL1122-05606 Fab or HL1162-65809 Fab, related to Figure 8. (A and B) Schematic showing the interactions between individual residues of (A) RL1122-05606 Fab (HC, red; LC, orange) or (B) HL1162-65809 Fab (HC, brown; LC, orange) and HA (green). Dotted lines represent polar interactions; inverted parentheses represent nonpolar interactions. The thick solid line circumscribes the interface essentially conserved between two Fabs. (C) Affinity measurement of HL1162-65809 against H3/KS. Kinetics of binding intensity was measured using BLI. Measured and fitted curves are shown in black and red, respectively. Blue line segregates the association (left side) and dissociation (right side) curves. Calculated K_D_ value and on/off-rates (k_a_ and k_d_) are shown below the graph. (D) Binding curves of IGHV14-4/IGKV4-59 mAbs against anti-mouse Igκ+λ capture Abs (left), K03.12 human IgG1 (middle), or humanized BMPC23 IgG1 (right). Each mAb including control (anti-human IgG-Fc polyclonal mouse Ab) was serially diluted from 1µg/ml and assessed in an ELISA. OD450 values at each dilution are plotted.

## Supplementary table titles and legends

**Table S1.** Summary results of single GC B cell cultures. The numbers of mice utilized for Nojima cultures and the numbers of cells sorted, recovered with IgG production, and exhibited HA-specificity are shown by each experimental condition. Although No Ab controls appeared independently in both low and high dose conditions of passive transfer experiments, the numbers of mice and cells were pooled in No Ab group.

| Transferred Ab | Days of harvest<br>(post-immunization) | Nos.<br>Mice | Nos.<br>Sorted cells | Nos.<br>IgG<sup>+</sup> cultures | Nos.<br>HA<sup>+</sup> Cultures |
| --- | --- | --- | --- | --- | --- |
| No Ab | 12 | 14 | 12864 | 1895 | 679 |
| HSV (20 µg) |  | 14 | 17856 | 3434 | 1147 |
| RBS (20 µg) |  | 14 | 26784 | 4845 | 1050 |
| HSV (200 µg) |  | 14 | 19678 | 3907 | 1442 |
| RBS (200 µg) |  | 14 | 26879 | 4769 | 1082 |
| No Ab | 16 | 16 | 14592 | 2486 | 1045 |
| HSV (20 µg) |  | 10 | 10944 | 2014 | 842 |
| RBS (20 µg) |  | 10 | 19104 | 3566 | 1138 |
| HSV (200 µg) |  | 12 | 14592 | 2962 | 1259 |
| RBS (200 µg) |  | 11 | 21119 | 3658 | 1087 |
| Total |  | 129 | 184412 | 33536 | 10771 |

**Table S2.**
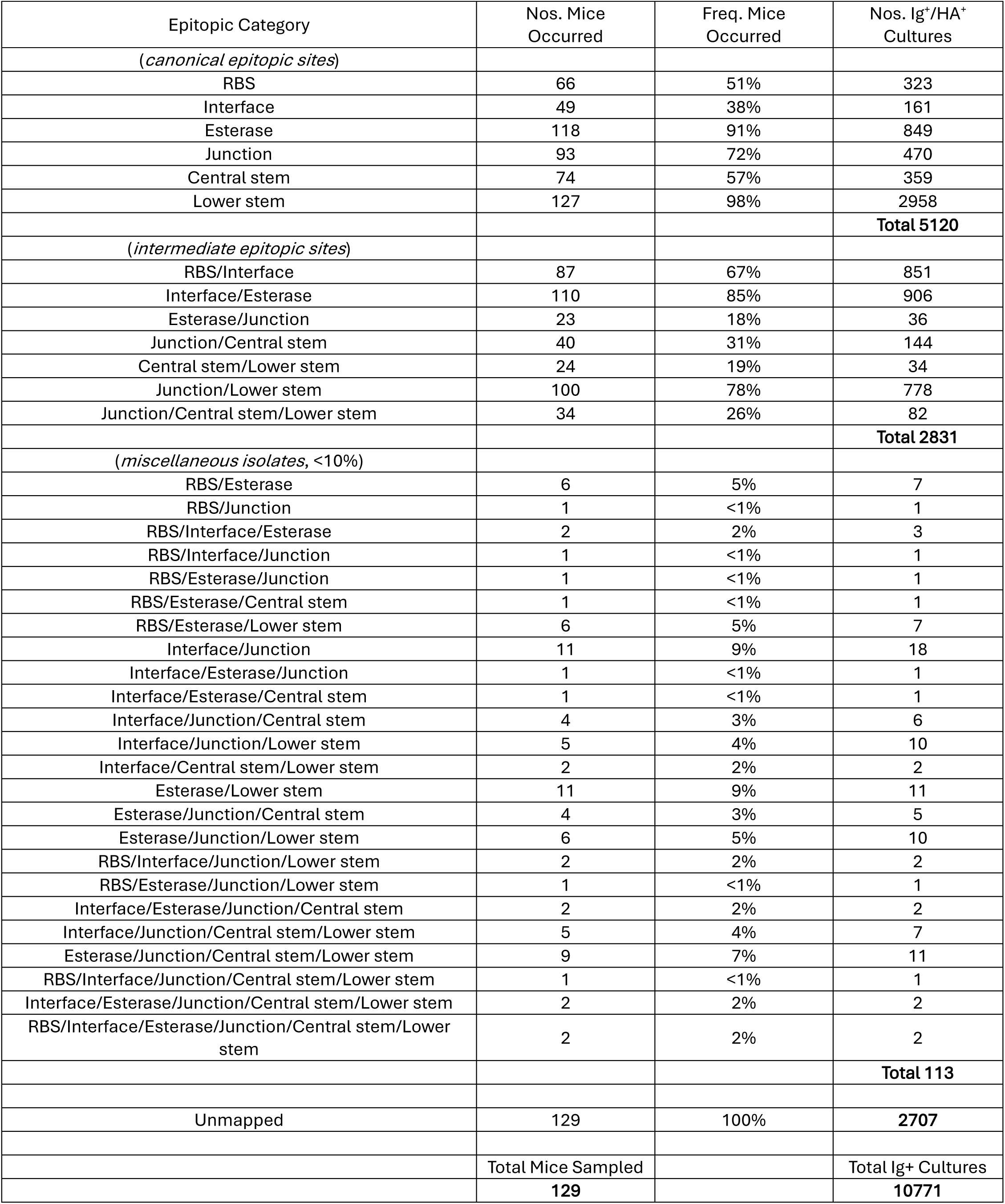
Summary results of epitope mapping in HA^+^ GC B cells. Table shows how often and how many each epitopic category was identified in mice from the entire experiments (n=129). If cells were identified in the categories which were observed in ≤ 12 mice (< 10% of 129 mice), they are pooled as miscellaneous isolates.

**Table S3.** Summary results of BCR gene sequencing. Numbers of cells recovered with complete heavy plus light chain pairs and unique clones identified from those sequences are shown. The numbers of mice and sequenced cells are same as those shown in Table S1. All of the HA^+^ cells were sequenced.

| Transferred Ab | Days of harvest<br>(post-immunization) | Nos.<br>Mice | Nos.<br>sequenced cells | Nos.<br>H+L chain pairs | Nos.<br>unique clones |
| --- | --- | --- | --- | --- | --- |
| No Ab | 12 | 14 | 679 | 451 | 267 |
| HSV (20 µg) |  | 14 | 1147 | 774 | 350 |
| RBS (20 µg) |  | 14 | 1050 | 773 | 360 |
| HSV (200 µg) |  | 14 | 1442 | 665 | 342 |
| RBS (200 µg) |  | 14 | 1082 | 603 | 256 |
| No Ab | 16 | 16 | 1045 | 806 | 363 |
| HSV (20 µg) |  | 10 | 842 | 596 | 239 |
| RBS (20 µg) |  | 10 | 1138 | 746 | 226 |
| HSV (200 µg) |  | 12 | 1259 | 733 | 266 |
| RBS (200 µg) |  | 11 | 1087 | 772 | 216 |
| Total |  | 129 | 10771 | 6919 | 2885 |

**Table S4.** Cryo-EM data collection, refinement and validation statistics.

|  | HL1162-65809 Fab<br>immunocomplex<br>PDB: 35VS | RL1122-05606 Fab<br>immunocomplex<br>PDB: 35WC |
| --- | --- | --- |
| **Data collection and processing** |  |  |
| Magnification | 165,000× | 165,000× |
| Voltage (kV) | 300 | 300 |
| Electron exposure (e<sup>-</sup>/Å<sup>2</sup>) | 49.7129 | 51.1227 |
| Defocus range (μm) | -0.8, -2.0 | -0.8, -2.0 |
| Pixel size (Å) | 0.74 | 0.74 |
| Symmetry imposed | C1 | C1 |
| Initial particle images (no.) | 1,171,321 | 4,684,294 |
| Final particle images (no.) | 256,207 | 587,026 |
| Map Resolution (Å) | 3.2 | 2.87 |
| FSC threshold | 0.143 | 0.143 |
| Map resolution range (Å) | 2.5 - 7.5 | 2.4 - 7.5 |
| **Refinement** |  |  |
| Initial model used | Modelangelo (*de novo*) | Modelangelo (*de novo*) |
| Map-sharpening B-factor (Å<sup>2</sup>) | -118.2 | -103.6 |
| Model composition |  |  |
| Non-hydrogen atoms | 7,427 | 7,402 |
| Protein residues | 953 | 951 |
| **Validation** |  |  |
| Clashscore | 1 | 1 |
| rotamer outliers (%) | 0.4 | 0.2 |
| Ramachandran plot |  |  |
| Favored (%) | 96.0 | 98.0 |
| Allowed (%) | 4.0 | 2.0 |
| Disallowed (%) | 0.0 | 0.0 |

